# Simple scaling laws control the genetic architectures of human complex traits

**DOI:** 10.1101/2022.10.04.509926

**Authors:** Yuval B. Simons, Hakhamanesh Mostafavi, Courtney J. Smith, Jonathan K. Pritchard, Guy Sella

## Abstract

Genome-wide association studies have revealed that the genetic architectures of complex traits vary widely, including in terms of the numbers, effect sizes, and allele frequencies of significant hits. However, at present we lack a principled way of understanding the similarities and differences among traits. Here, we describe a probabilistic model that combines mutation, drift, and stabilizing selection at individual sites with a genome-scale model of phenotypic variation. In this model, the architecture of a trait arises from the distribution of selection coefficients of mutations and from two scaling parameters. We fit this model for 95 diverse, highly polygenic quantitative traits from the UK Biobank. Notably, we infer similar distributions of selection coefficients across all these traits. This shared distribution implies that differences in architectures of highly polygenic traits arise mainly from the two scaling parameters: the mutational target size and heritability per site, which vary by orders of magnitude across traits. When these two scale factors are accounted for, the architectures of all 95 traits are nearly identical.

## Introduction

A central goal of genetics is to understand how genetic variation maps to phenotypic variation. Starting in the late 20th century, there was huge progress toward identifying the genes for Mendelian traits. But most phenotypic variation in humans is genetically complex, and it is only in the last 15 years that genome-wide association studies (GWAS) have started to reveal the genetic basis of variation in a wide array of complex traits [1]. These studies have now identified tens of thousands of robust associations between genetic variants and a wide array of traits and diseases.

One intriguing observation from this work is the striking variation in genetic architecture among complex traits [2–5]. (Here, we use the term *architecture* to refer to the numbers of causal variants and their joint distribution of allele frequencies and effect sizes.) Traits have been found to vary in all aspects of genetic architecture, including: the number and magnitude of significant signals found at a given sample size [6]; the fraction of heritability explained by lead GWAS signals [3]; the allele frequency distributions of significant variants [4, 7]; the estimated numbers of causal variants [8, 9]; and the SNP-based heritability [10, 11].

Nonetheless, diverse traits do show important similarities. First, most complex traits are influenced by large numbers of variants with small effects, only a small fraction of which can be confidently detected at current sample sizes [12–14]. Indeed, even relatively “simple” complex traits such as molecular biomarkers are highly polygenic with ∼10^4^ causal variants spread widely across the genome, compared to ∼10^5^ or more variants for traits such as height or BMI [2, 9, 15–17].

Second, the distributions of effect sizes of causal variants are not fit well using standard modeling assumptions such as normal distributions. Instead, effect sizes typically span several orders of magnitude, much like power-law distributions [18, 19].

Third, trait-associated variants are often highly pleiotropic: i.e., they influence many traits simultaneously. Many pairs of traits show significant genetic correlations, indicating that allelic impacts are often shared [16, 20, 21]; moreover, whenever different traits are mediated through overlapping cell types or pathways, we can expect that they will share many of the same regulatory variants even if the directions of effects are uncorrelated [5, 14, 16, 22].

Fourth, selection plays a central role in shaping complex trait architecture. Evolutionary theory predicts that variants with phenotypic effects would usually be under selection and, in particular, that selection is usually stronger for larger-effect variants [23]. Consistent with this, variants with larger effect sizes tend to be at lower frequencies, indicating that selection prevents such variants from reaching high frequencies [24–26]. Since heritability depends on both effect sizes and allele frequencies, an important consequence is that the genes that are most important for a trait contribute less to heritability than would be expected in the absence of selection, thus flattening the heritability distribution across genes [17, 27, 28].

Here we develop a principled approach for understanding similarities and differences in genetic architecture. Specifically, we want to understand how the population genetic processes of mutation, selection, and drift alongside properties of individual traits determine the numbers of variants, as well as the joint distributions of allele frequencies and effect sizes. *What features of these processes are shared across traits? And which are different?*

To answer these questions, we require a model for how population genetic processes shape complex trait architecture. Current models differ primarily in their assumptions about the relationship between selection on alleles (or alternatively their frequencies), and the effect sizes of those alleles on a trait of interest [23]. The heuristic ‘*α*-model’, developed for estimating SNP heritability, assumes a particular parametric relationship between allele frequencies and effect sizes [25, 29, 30]. While the *α*-model is motivated by the observed inverse relationship between effect size and frequency, the precise functional form is arbitrary. In turn, several evolutionary models postulate particular parametric relationships between the strength of selection on alleles and their effect sizes, and then rely on explicit population genetics models to derive the relationship between allele frequencies and effect sizes and other aspects of genetic architecture [31–36]. These models, however, differ in their predictions about architecture, owing to the various ad-hoc parametric relationships they assume.

Simons et al. (2018) introduced an evolutionary model that moves beyond ad-hoc choices by deriving the relationship between selection on alleles and their effects on a trait under an explicit, interpretable, biological model [27]. Motivated by extensive evidence that many quantitative traits are subject to stabilizing selection, where fitness declines with displacement from an optimal trait value [23, 37, 38], and that genetic variation affecting one trait often affects many others [5, 16, 22], they modeled selection on alleles that arises from stabilizing selection in a multi-dimensional trait space. They then used an explicit population genetic model to derive the genetic architecture of a focal trait with mutation, genetic drift, and stabilizing selection in a multi-dimensional trait space.

As we will show in the next section, the Simons et al. model can be reframed as a generative (statistical) model for the genetic architecture of a continuous complex trait, which depends on a few biologically interpretable parameters. We next describe approaches to infer these parameters and test the model fit based on data from GWAS. Applying our inference to 95 diverse, highly-polygenic quantitative traits from UK Biobank we show that this model provides an excellent fit to the data. Surprisingly, we find that most variation in architecture among traits is explained by differences in just two scaling parameters: the mutational target size and heritability per site.

## Results

### A population genetic model of complex traits

As a starting point, we assume that phenotypic variation exists in a high-dimensional trait space under stabilizing selection. Here we outline key elements of the model; further details and biological motivation can be found in the Supplement and in [23, 27].

We model each person’s phenotype as a point in an *n*-dimensional trait space, and assume that this dimension is high (*n* ≥ 10). To model stabilizing selection we assume that there is an optimal phenotype, and that fitness decreases with Euclidean distance from the optimum (Figure 1A).

**Figure 1:**
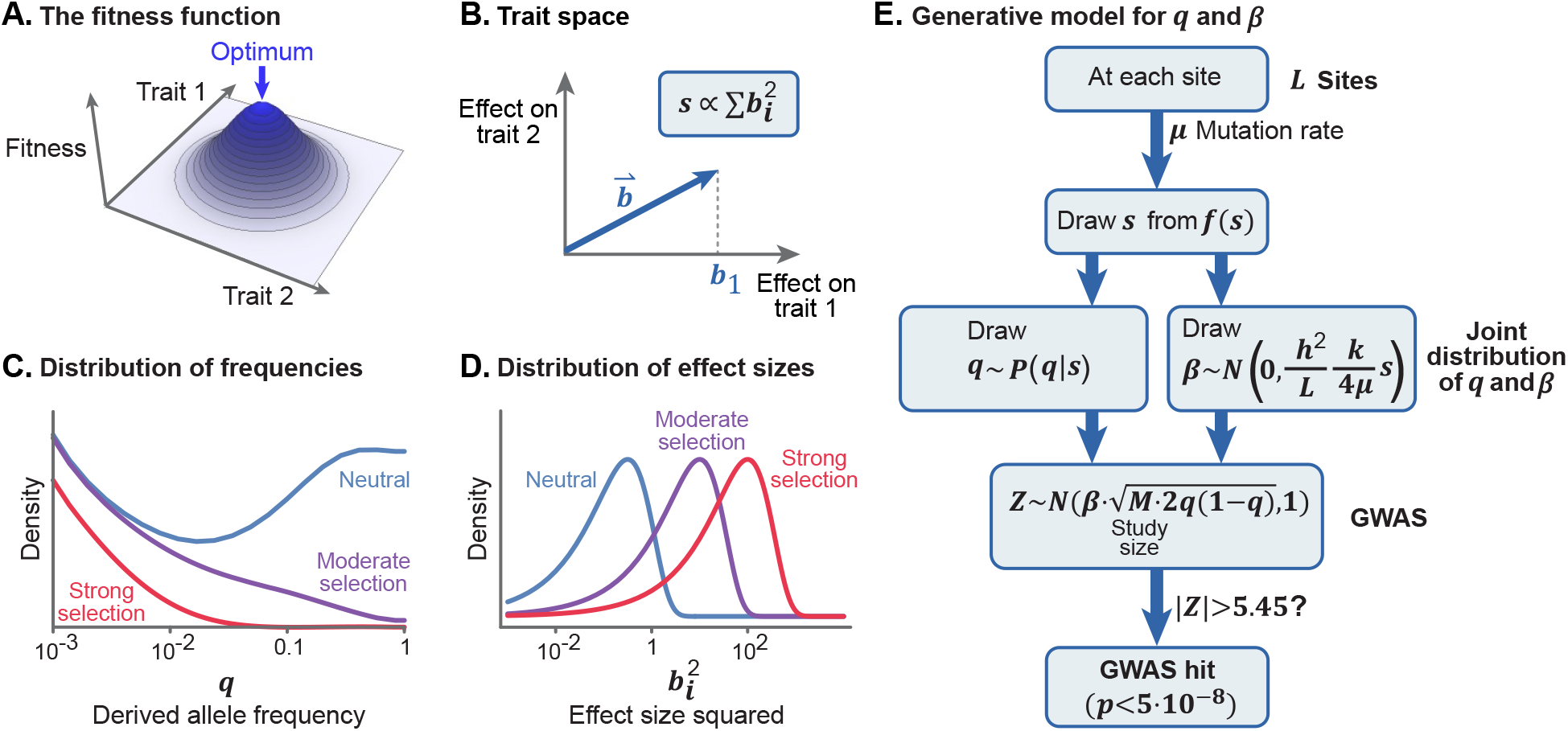
The model. **(A)** We use Fisher’s concept of a multi-dimensional trait space. Under stabilizing selection, an individual’s fitness declines with distance from the optimal phenotype [40]. **(B)** The selection coefficient experienced by a variant is proportional to the sum of squared effects on all traits. **(C)** We compute the distribution of derived allele frequencies (q) conditional on s and demography. **(D)** The distribution of effect sizes for Trait 1 (b_1_) is normally distributed given s. **(E)** The generative model for q, β, and the observed Z-score at any given site.

The phenotypic effect of each variant is represented by a random vector in the *n*-dimensional trait space, namely 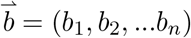, where *b*_*i*_ is the additive effect of the derived allele on the *i*-th trait. We assume that a person’s phenotype arises from their genotype according to the standard additive model in *n*-dimensions: it is a vector sum over the effects of all variants plus a random vector representing the environmental effects [39].

The model thus far is mathematically similar to Fisher’s Geometric model [40], which Fisher and others used to study adaptive processes [41], but we consider a different question and a different evolutionary setting. We focus on the genetic architecture of a single highly polygenic trait that arises in the balance between mutation, stabilizing selection in the multidimensional trait space, and genetic drift.

#### Mutation, selection, and drift at individual sites

Each generation, mutation introduces new trait-affecting variants into the population at a rate *μ* per site, per gamete, per generation. The long-term fate of variants is determined by the combined action of selection and drift.

Under stabilizing selection, at equilibrium, selection holds the phenotypic mean very close to the optimal phenotype, and thus acts against mutations (and against variation in general). The strength of selection, *s*, acting against a variant is proportional to its squared magnitude in the *n*-dimensional trait space (Figure 1B):

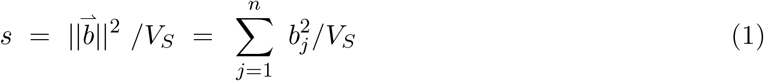

with *V*_*S*_ reflecting the width of the fitness function around the optimum.

Given *s* we can compute the present day allele frequency distribution, as follows. When traits are subject to stabilizing selection, selection at individual sites is under-dominant, meaning that selection acts against minor alleles, regardless of the direction of effect [27,42,43]. At strongly selected sites, this approximates the standard model of selection against deleterious alleles. Specifically, the expected change in allele frequency at an autosomal site in a single generation, given current derived allele frequency *q* is

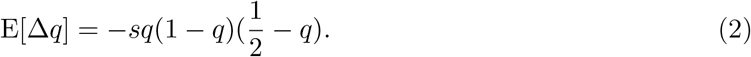

Meanwhile, the variance in the change in allele frequencies, i.e., drift, scales inversely with population size. Hence, the distribution of present-day allele frequencies is the result of a stochastic process including past mutations, selection, and drift – which depends on the history of population sizes. For our analysis here, we computed the distribution of present day allele frequencies under the stabilizing selection model using a demographic model estimated for the British population [44]. As expected, strongly selected variants (large *s*) tend to be rare, while nearly-neutral variants (small *s*) can drift to high frequencies (Figure 1C).

#### The relationship between selection and effect sizes

Next we need to understand how selection in the multi-dimensional trait space relates to the genetic architecture of a single focal trait of interest. Without loss of generality, we focus on the first dimension in the *n*-dimensional trait space, and to simplify the notation we denote the effect size *b*_1_ of a variant on trait 1 simply as *b*.

While it seems natural to think about *s* as a function of a variant’s effects on traits, here we invert this relationship: specifically, we need the conditional distribution of the effect size on trait 1 given *s*. This conditional distribution reflects uncertainty about the projection of 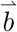 onto the first dimension if all we know is *s* (or equivalently 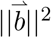). Fortunately, when the number of traits is sufficiently large, this conditional distribution is well approximated by a simple form (see [27] and SI Section 1.3), namely:

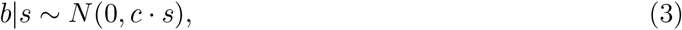

where *c* = *V*_*S*_*/n*. Intuitively, variants under weak selection (small *s*) tend to have small squared effect sizes (*b*^2^) and variants under strong selection (large *s*) tend to have larger squared effect sizes (Figure 1D).

With the distributions for *b* and *q* given *s*, we can now compute the expected per site contribution to phenotypic variance as a function of *s*, given by E[2*b*^2^*q*(1 − *q*)|*s*]. Under non-equilibrium demography, the expectation does not have a simple form but it is plotted in Figure S2. At sites where selection is weak, *b*^2^ is small, and these sites contribute little to the overall genetic variance, *V*_*G*_. When selection is strong, *b*^2^ is large, but selection holds *q*(1 − *q*) low, and these effects cancel out, so these sites are capped in terms of how much they can contribute to *V*_*G*_ [27].

#### Single-site dynamics and heritability

Moving from single sites to a genome-wide model, let *L* be the number of sites in the genome at which mutations can affect Trait 1; we refer to *L* as the *mutational target size*. We use *f* (*s*) to denote the unknown distribution of selection coefficients of mutations at these *L* sites. Then the (expected) total additive genetic variance for Trait 1 is given by

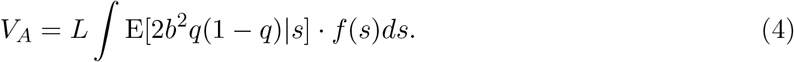

Next, we rescale *b* from the original but arbitrary measurement units into units of standard deviations of the trait value: we define 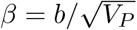, where *V*_*P*_ is the phenotypic variance. Dividing both sides by *V*_*P*_, and noting that *V*_*A*_*/V*_*P*_ is the (narrow sense) heritability *h*^2^, we can relate heritability to the site-level parameters:

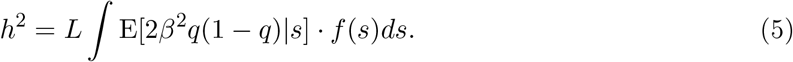

This equation expresses the key relationship between heritability (*h*^2^), mutational target size (*L*), and the expected contribution to variance per site.

Finally, Equation 3 can be rewritten in terms of *β* and population genetics parameters (for details see SI Section 1.5):

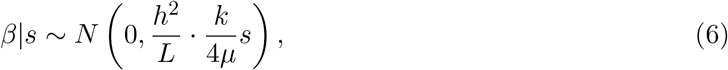

where *k* is a constant that depends on *f* (*s*) and demography and is approximately 1, and *μ* is the mutation rate. Crucially, Equation 6 shows that the trait’s heritability per site, *h*^2^*/L*, is a fundamental scaling factor that relates selection on alleles to their effects on the trait.

Together, these results provide a generative model for the genetic architecture of a complex trait (Figure 1E). Assuming that the demographic history and mutation rate per site are known in advance, this model is fully specified in terms of three unknowns: the mutational target size, *L*; the heritability per site, *h*^2^*/L*; and the distribution of selection coefficients, *f* (*s*). We now describe how we estimate these from GWAS data.

### Inference of model parameters from GWAS data

In principle we would want to perform inference using all causal variants, but this is technically challenging since most causal sites have very small effect sizes; hence there is great uncertainty about which sites are causal and their true effect sizes. As a tractable alternative, we restricted our inference to the independent genome-wide significant hits for each trait. We account for this restriction in the inference by noting that we only observe the subset of sites for which the absolute GWAS z-score exceeds 5.45, corresponding to the conventional significance threshold of *p<*5*×*10^−8^.

We performed simulations to illustrate the changes in architecture at GWAS hits as a function of each of the main model components (Figure 2; SI Section 5). As expected from theory:

**Figure 2:**
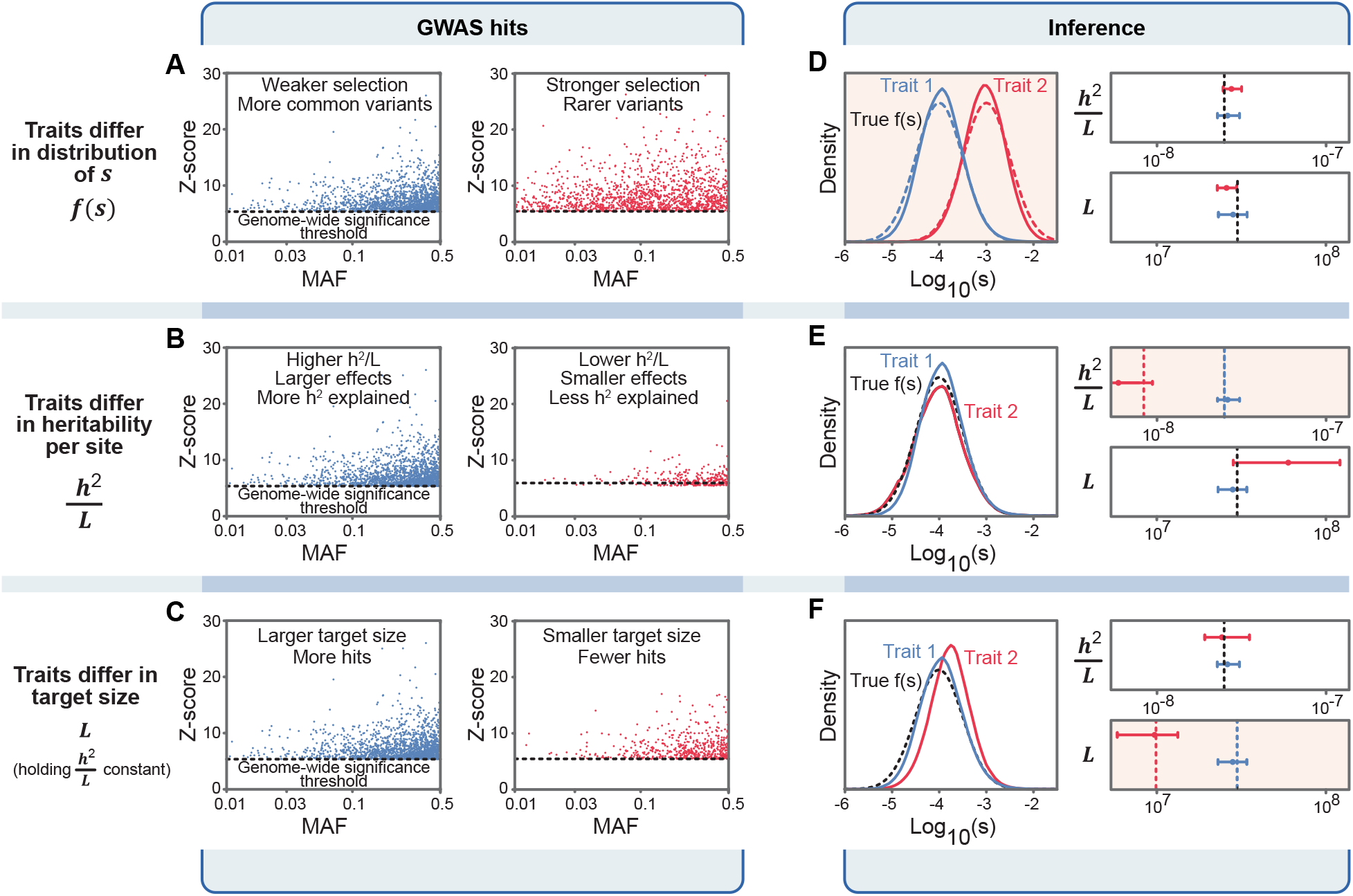
Inference of model parameters. **(A)–(C)** The joint distributions of minor allele frequencies (MAFs), z-scores, and numbers of hits per trait depend on model parameters as illustrated here. Each graph shows simulated distributions of genome-wide significant hits, with the graphs in each row differing in one of the main axes of our model. **(D)–(F)** True values of the distributions of f (s) as well as h^2^/L and L are indicated by the dotted lines; inferences are indicated by solid lines or by point estimates with bars indicating 90% bootstrap CIs.

A. When selection is weak, causal variants can drift to high frequencies, and most significant hits are at common variants. Conversely, when selection is strong, there is a greater fraction of rare variants among the significant hits, and an inverse relationship between effect size and allele frequency.
B. When traits have high heritability per site *h*^2^*/L*, the squared effect sizes and z-scores tend to be larger, there are more genome-wide significant hits and they explain a greater proportion of heritability, compared to traits with low *h*^2^*/L*.
C. When traits have a large mutational target size *L* (holding *h*^2^*/L* and *f* (*s*) constant), there are more causal variants, and more genome-wide significant hits, but the distribution of allele frequencies and effect sizes, and the proportion of heritability explained by hits, are unaffected.

We implemented a maximum likelihood method that estimates the components of our model from the joint distribution of *q* and |*z*| across significant hits (Figure 2D-F; SI Section 5). We fit *f* (*s*) using a spline function with four knots, thus our full model includes six parameters per trait: four for *f* (*s*), as well as *h*^2^*/L*, and *L*.

We tested this method using simulated GWAS data under a variety of parameter values. We find that even with modest numbers of hits (∼100) the method provides accurate estimates, while the estimates are noisy for traits with fewer hits (e.g., panel 2E). It may seem surprising that we can estimate *f* (*s*) from relatively few observations, but each variant carries considerable information about the strength of selection: the allele frequency bounds *s* for that variant from above and the effect size bounds *s* both from below and above, such that jointly they are highly informative (Figure S3). Given these results we analyzed traits with at least 100 hits.

We also observed that the data are less informative at both ends of the range of possible selection coefficients: GWAS has low power to detect very strongly selected variants (*s* ≳ 10^−2^) as the allele frequencies are too low, and low power to detect effectively-neutral variants (*s* ≲ 10^−5^) as their effect sizes are too small. We therefore implemented a regularization penalty to constrain *f* (*s*) to sensible values at extremes of the range (SI Section 3.8).

### Dataset of 95 quantitative traits from the UK BioBank

We selected traits from the UK Biobank for analysis, as follows (SI Section 2.4). Since our model is most directly applicable to quantitative continuous traits, we restricted our analysis to such traits. We identified independent lead variants for each trait using COJO [45]. Since low-frequency variants are often poorly imputed, we removed hits with MAF<1% (see SI Section 6.3 for how we account for this in the inference). We excluded traits for which more than 10% of hits fell within a single LD block, as well as hits in regions of extremely high LD (LD score *>*300). As noted above, we restricted ourselves to traits with at least 100 independent hits; doing so implies that the traits are among the more polygenic and heritable traits in UKBB. For each trait we recorded the number of hits, and the estimated allele frequency, z-score, and effective sample size for each hit.

This resulted in a list of 95 traits that passed all filtering steps, with a range of 100 to 1,426 hits per trait (mean=398). These traits include 40 morphometric traits, of which 26 are related to body weight or adiposity (e.g., BMI, waist circumference and birth weight) as well as 14 others (e.g., height, bone mineral density and hand grip strength). The traits also include 27 blood phenotypes (e.g. platelet traits, lymphocyte count, and hemoglobin measurements), and 12 molecular traits sampled from blood or urine (e.g., IGF-1, triglycerides and calcium levels). Additionally, we have 9 cardiovascular traits, including pulse rate, blood pressure measurements, and pulmonary function traits. Lastly, we include 6 ophthalmologic traits, and 1 behavioral trait (age at first sexual intercourse).

### Distributions of trait parameters

We applied our inference to all 95 traits. Figure 3A shows the estimated distributions of selection coefficients, *f* (*s*), for five traits that vary in polygenicity and kind, from calcium levels to BMI. Note that although our inference is based on significant hits, *f* (*s*) represents the distribution of *s* among *new* mutations.

**Figure 3:**
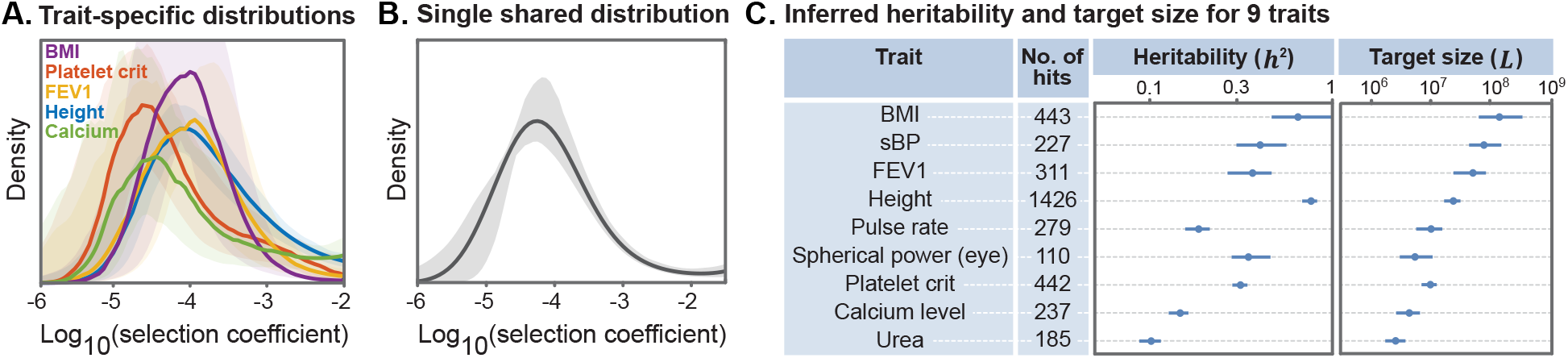
Parameter estimates for example traits. **(A)** Distributions of selection coefficients, f (s), estimated for each trait separately, using trait-specific distributions, with 90% confidence envelopes. **(B)** Single shared distribution (SSD) of selection coefficients, estimated using all 95 traits together. **(C)** Properties of example traits. h^2^ and L are estimated using f(s) from the SSD, which helps to stabilize the estimates.

We infer that most trait-affecting mutations are under weak selection, with *s* ranging between 10^−5^ and 10^−3^. In this range, the strength of selection is roughly comparable to genetic drift (∼ 10^−4^ per generation), consistent with the observation that many GWAS variants are common, and more generally that much of the heritable variation in complex traits arises from common variants [3, 4, 10, 12]. However, the distributions have a substantial tail in the strong selection range (*s >* 10^−3^), and therefore span multiple orders of magnitude. Since selection coefficients span multiple orders of magnitude so too, should effect sizes. This echoes recent results showing that distributions of effect sizes for complex traits do indeed span multiple orders of magnitude (in contrast to the normal distribution which has often been assumed in statistical genetics models) [18, 19].

Importantly, the distribution of selection coefficients, *f* (*s*), is similar across all 95 traits, with the confidence envelopes for different traits largely overlapping (Figures 3A, S5). We thus conjectured that we could build a unified model by assuming a Single Shared Distribution for *f* (*s*), which we refer to as the SSD, instead of assuming separate Trait-Specific Distributions (TSDs). The SSD is shown in Figure 3B. As we will show, the SSD provides a useful approximation for the architecture of individual traits, while greatly cutting down the number of model parameters and highlighting important shared features of trait architecture.

In contrast to *f* (*s*), the heritability *h*^2^ and especially the mutational target size *L* vary markedly among traits. For example, among the traits in Figure 3C, BMI (444 hits) has an estimated target size of 150 MB, one of the highest estimated target sizes, in contrast to urea (185 hits) with a target size of 2.5 MB, one of the lowest estimated target sizes. These results are broadly consistent with expectations from previous studies of polygenicity showing that morphological traits including height and BMI have many more contributing variants than do molecular traits, illustrated here by urea and calcium [2, 9]. Our estimates of heritability vary less widely among traits, and are concordant with previous estimates (Table S1; [10]).

### Quantifying model fit

Next, we assessed the fit of our models to the genetic architecture observed in GWAS, including how the fit is affected by using the SSD approximation (SI Section 4). To do so, we computed a measure of model fit using the predicted distribution of z-scores given the allele frequencies and study size. For each variant *i*, we computed what we refer to as a *residual p-value*: Pr(|*z*| *> z*_*i*_ | *z*_*i*_ *>* 5.45, *q*_*i*_, *Model*), where *z*_*i*_ and *q*_*i*_ are the observed z-score and frequency of SNP *i*, respectively, and *Model* indicates the SSD or TSD model. The SSD model fit depends on only one trait-specific parameter, *h*^2^*/L*, as *f* (*s*) is shared across traits, and the residual p-value statistic is independent of *L*. In contrast, the TSD model fit depends on five parameters per trait: *h*^2^*/L* and four parameters to fit *f* (*s*).

The residual p-value has a simple interpretation: If we correctly model the distribution of z-scores among significant hits, then the distribution of residual p-values will be uniform between 0 and 1. If the observed z-scores are too small then the residual p-values will skew toward 1, and if the z-scores are too high they will skew toward 0. To avoid overfitting, we split the genome into approximately independent blocks [46], each time inferring the model on 90% of the blocks and computing residual p-values for the held-out 10%.

We first considered the fit of our model for height, the trait with the greatest number of hits in our dataset (Figure 4A). For height, the distribution of z-scores across the 1426 hits is fit essentially perfectly by the SSD model (Figure 4A). In contrast, two simpler heuristic models provide a poor fit to the distribution of z-scores (Figure 4A). First, when we assumed that effect sizes are normally distributed, the resulting residual p-values deviate greatly from uniformity with many extremely small p-values. We also considered a version of the *α*-model with a normal density of effect sizes conditional on allele frequencies [30]. By fitting the inverse relationship between allele frequencies and effect sizes, the alpha model improves the overall fit, but still has an excess of tiny p-values. In both cases, the underlying normal distribution is too narrow to accommodate the wide variation in observed effect sizes, including many hits close to the significance threshold and a minority of much stronger hits.

**Figure 4:**
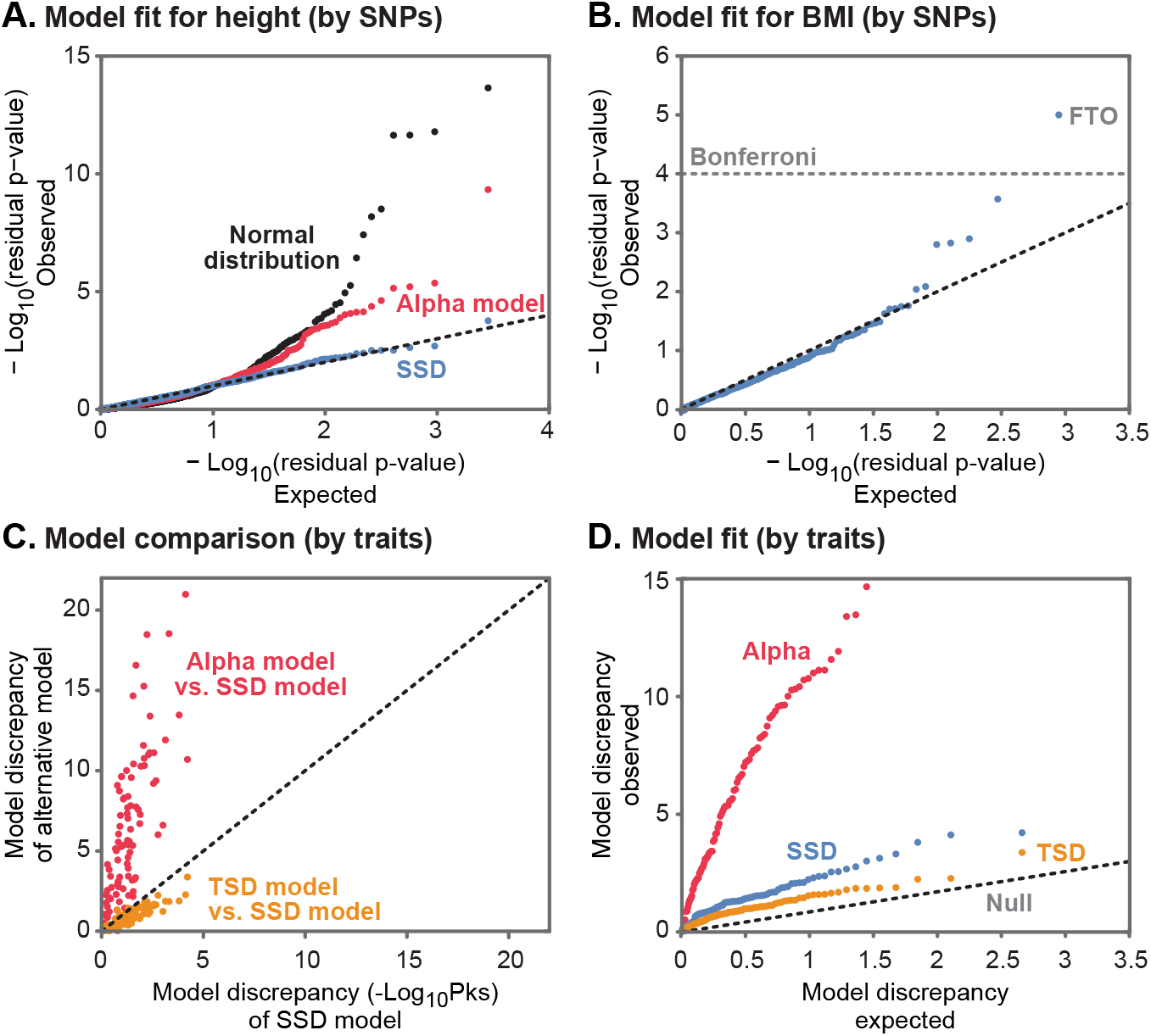
Model fit. **(A)** QQ-plot of residual p-values for height (each data point is a SNP) under three models: the SSD model provides a good fit to the distribution of z-scores, while two other models fit poorly. **(B)** For BMI, the SSD model fits most of the z-score distribution, but a few hits are more significant than expected, notably at FTO. **(C)** Goodness-of-fit comparison among models (each data point is a trait). Red: α-model versus SSD shows much worse fit for the α-model (data above diagonal). Orange: TSD versus SSD generally shows slightly better performance for TSD (data below diagonal), while using five parameters per trait instead of one. **(D)** QQ-plots for model fits (by trait) for TSD, SSD, and α-models.

For BMI, with 444 hits, our SSD model fits most of the distribution well, but the top hits are larger than expected (Figure 4B). In particular, the residual p-value of one SNP is significant even after Bonferroni correction. Unsurprisingly, this outlier represents the well-known FTO signal that was detected even in very early GWAS studies [47, 48], and that also appears as an outlier for 14 other morphometric traits in our dataset.

We find 12 additional outlier SNPs for a variety of other traits. The 12 outliers include both missense and noncoding variants, and are all found near strong candidate genes for the relevant traits (Table S2). We hypothesize that these outliers violate our model assumptions in some way that allows them to be common despite having a large effects. For example, they might have much smaller pleiotropic effects than most other variants affecting those traits, leading to weaker selection than expected given their effect sizes. Alternatively, they may have been targets of strong positive or balancing selection that allowed them to reach high frequencies despite their large effect sizes.

We next performed goodness-of-fit tests for each trait to determine whether the overall distributions of residual p-values match the expected uniform distribution, using Kolmogorov-Smirnov statistics (Figure 4C; SI Section 4). For most traits, the TSD model fits slightly better than the SSD model, indicating that the TSD correctly identifies some degree of trait-specific signal, though it uses five parameters to do so, compared to one, for SSD. In contrast, the *α*-model fits the data far worse than the SSD model. Similarly, Figure 4D shows QQ-plots of model-fits across models and traits. Both the SSD and TSD models show a modest inflation of p-values, perhaps relating to simplifying assumptions in the model; nonetheless, both models fit the data well, as we can only reject these models for 1, and 4 traits out of 95, for the TSD and SSD models, respectively. Thus, we conclude that the SSD model provides an accurate, yet parsimonious, description of genetic architecture.

### Prediction of allele ages

The previous results show that the model provides a good empirical fit to the data. We next wanted to evaluate whether it can also predict the evolutionary processes underlying the genetic architecture. To do this, we turned to an entirely different type of predictions from our model. Conditional on allele frequency, deleterious alleles tend to be younger than neutral alleles [24,49]; our parameter estimates can be used to predict the extent of this effect for GWAS hits.

We compared our model’s predictions to allele ages estimated from a reconstruction of the ancestral recombination graph using Relate [44] (see SI Section 7 for details, including bias correction).

Looking at the allele ages of GWAS hits for all 95 traits as estimated by Relate, we see that they are much younger than frequency-matched, putatively neutral alleles (Figure 5). We estimate that the median age of a GWAS hit variant is 173,000 years, compared to 597,000 years for matched neutral variants. These observations highlight the competing influences of selection and drift on GWAS hits: due to selection GWAS hits are much younger than matched neutral variants, yet at the same time selection is weak enough that most of GWAS hits are fairly old, predating the out-of-Africa bottleneck. This finding echoes previous ones showing that many common GWAS hits are shared among African and non-African populations [50, 51].

**Figure 5:**
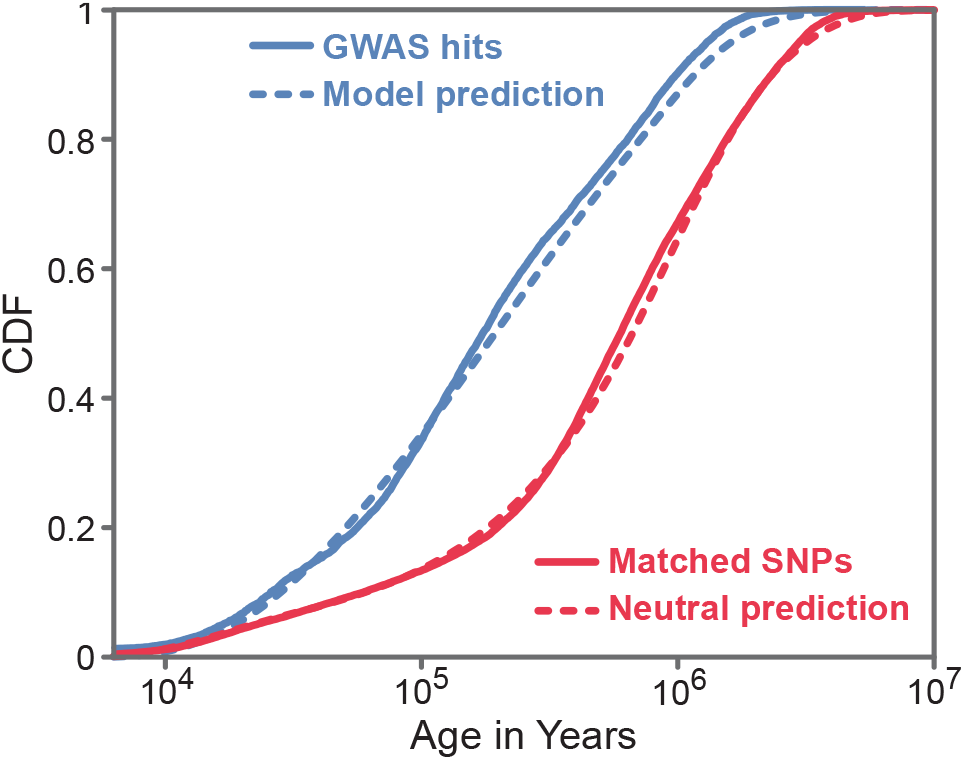
Allele ages. The distribution of GWAS hit allele ages for all 95 traits (solid blue), estimated using Relate, compared to the distribution of allele ages predicted by our model (dashed blue). Also shown, the distribution of allele ages for neutral frequency-matched SNPs (solid red) and the distribution predicted by a neutral model (dashed red). Allele ages were converted to years by assuming 28 years per generation [52].

Most importantly, the shift in ages of GWAS hits compared to matched control SNPs is predicted well by the SSD model, indicating that this model captures the right magnitude of selection coefficients. (For results about our ability to distinguish selection coefficients with this analysis see Figure S14.) Since the allele ages inferred by Relate are estimated from local haplotype structure, information that isn’t used by our inference, this concordance provides an external validation of the SSD model.

### Simple scaling rules control differences in trait architectures

The fit of the SSD model suggests an intriguing prediction: that the differences in genetic architecture among traits are primarily due to just two trait-specific parameters: the heritability per site (*h*^2^*/L*) and the mutational target size (*L*).

To see why, consider the full genetic architecture for a trait, for *all* variants regardless of whether they can be detected by GWAS. First, for a given demographic history, the distribution of allele frequencies depends only on the distribution of selection coefficients, *f* (*s*). Hence, under the SSD approximation, the allele frequency distribution is shared across traits and can be predicted from our estimate of *f* (*s*) (black line, Figure 6A).

**Figure 6:**
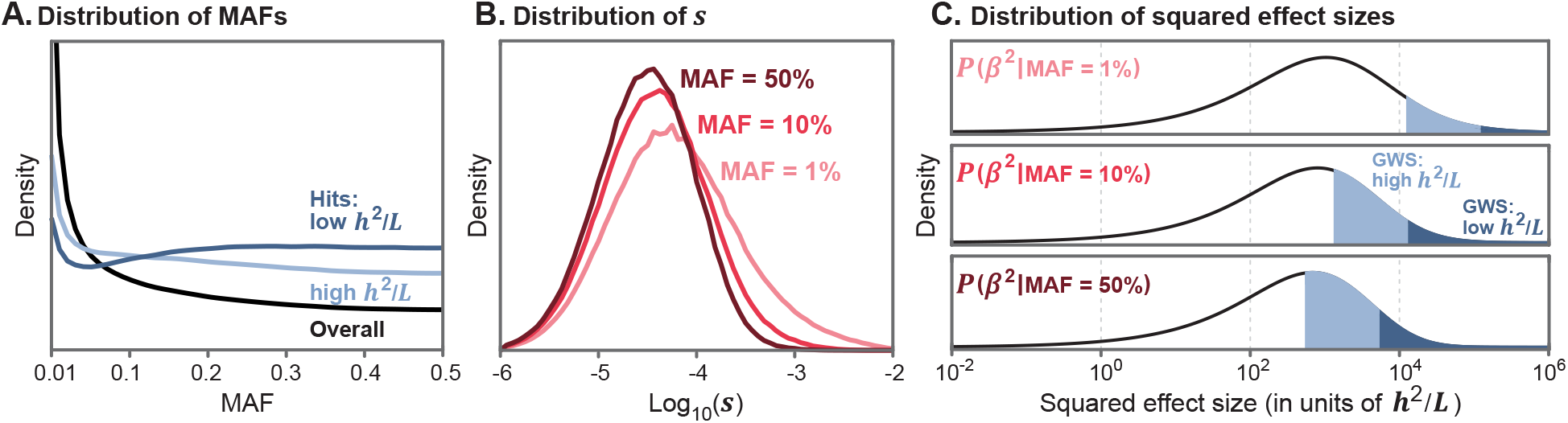
Shared genetic architecture under the SSD model. **(A)** Distribution of allele frequencies for all causal variants (black), and for genome-wide significant hits (light/dark blue), for our inferred f (s) and British population history. **(B)** Distributions of selection coefficients at causal variants with different minor allele frequencies. **(C)** Distributions of squared effect sizes β^2^, shown here for three example minor allele frequencies; notice that effect sizes are scaled by the natural units of h^2^/L. For traits with high h^2^/L, variants within both the light and dark blue regions are genome-wide significant (GWS); for traits with low h^2^/L, only the dark blue regions are significant.

Next, while *f* (*s*) represents the distribution of selection coefficients among new mutations, strongly selected variants are less likely to reach high frequencies. Hence, the distribution of selection coefficients shifts towards the left with increasing allele frequencies, as shown in Figure 6B.

Given the distributions of selection coefficients at different allele frequencies from Figure 6B, we can compute the distribution of squared effect sizes as a function of allele frequencies, by integrating Equation 6 over *s*. Crucially, under the SSD model, these distributions are identical across traits, if the effect sizes on the x-axis are scaled in terms of *h*^2^*/L* (black lines, Figure 6C).

How do these distributions affect GWAS hits? Unlike the underlying distributions, the power to detect significant variants in GWAS depends on the actual squared effect sizes *β*^2^, not scaled by *h*^2^*/L* (and it depends on allele frequency and sample size). Consequently, there is more power to detect variants for traits with higher *h*^2^*/L* – this is intuitive, because higher *h*^2^*/L* implies that each site explains more variance in the trait. This is illustrated in Figure 6C: for traits with high *h*^2^*/L*, variants within both light and dark blue regions are genome-wide significant, but for traits with low *h*^2^*/L*, only variants within the dark blue regions are detected.

Moreover, even though the underlying distribution of causal variant allele frequencies is shared among traits, the frequency distribution of hits is predicted to vary. For traits with high *h*^2^*/L* there is relatively more power to detect low frequency variants than for traits with low *h*^2^*/L* (Figure 6A).

Lastly, the second scaling parameter, *L*, represents the mutational target size. Conditional on *h*^2^*/L*, changing *L* only changes the *numbers* of causal variants (and numbers of hits) but does not change any of the distributions.

We first tested these predictions for three traits: height, platelet crit (a blood phenotype), and FEV1 (a measure of lung function), where we reduced the sample size to 330,000 so it is identical for all traits (SI Section 6.6). Figure 7A shows height and platelet crit. These two traits differ greatly in their number of hits (1245 vs 499), estimated heritability (76% vs 31%) and mutational target size (25 MB vs 10 MB). However, we estimate that they have very similar values of *h*^2^*/L* (∼3.2 *×* 10^−8^). Consistent with our model, the distributions of *z*-scores, effect sizes, and MAFs for significant hits are nearly identical for the two traits.

**Figure 7:**
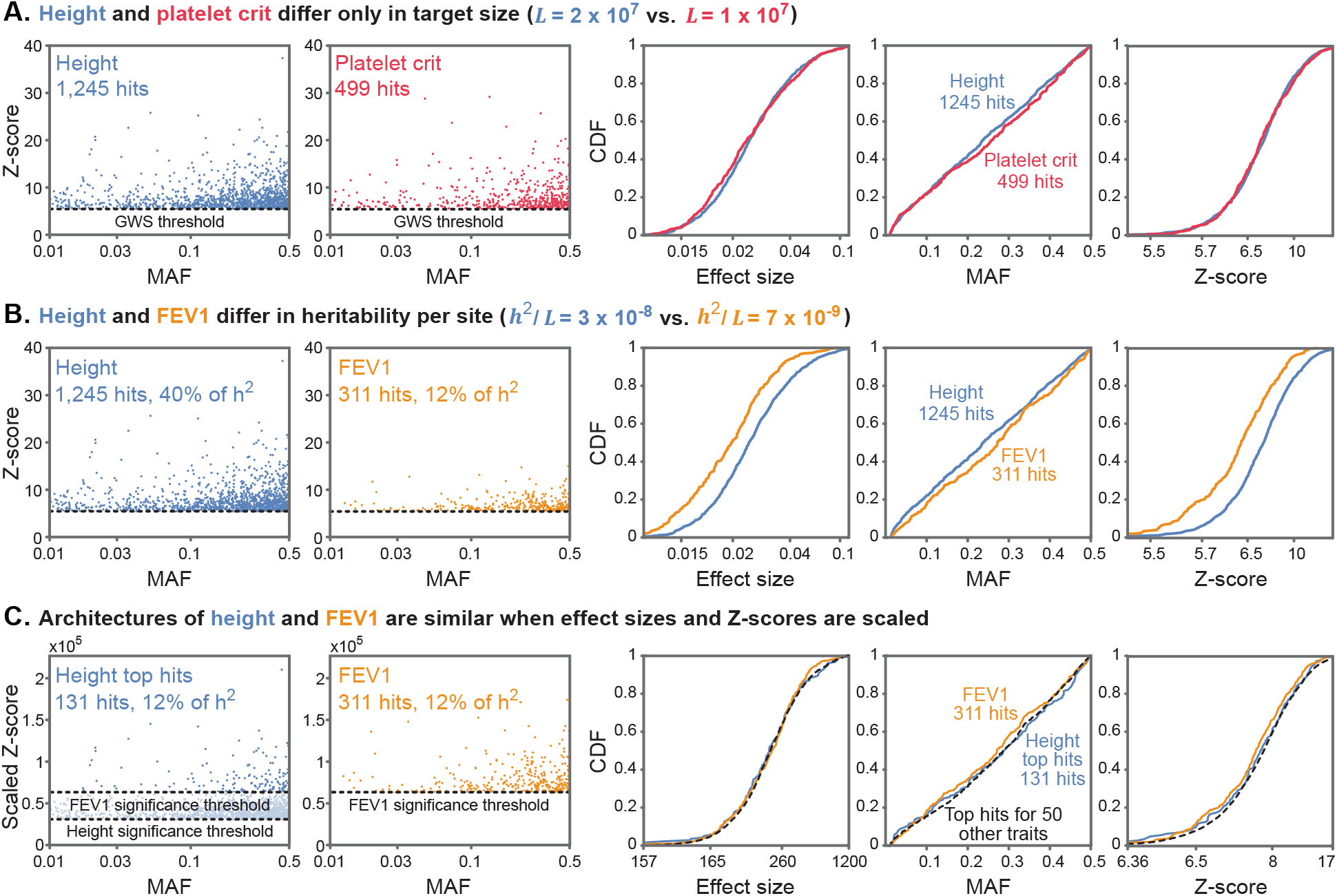
Heritability and target size underlie differences between trait architectures: examples for three traits. **(A)** Height (blue) and platelet crit (red) have the same heritability per site h^2^/L, but height has a much higher mutational target size L. This results in many more hits for height (1245) than for platelet crit (499) (2 left panels). However, the marginal distributions of z-scores, effect sizes, and MAFs of hits are nearly identical for the two traits (3 right panels). **(B)** Height (blue) and FEV1 (gold) differ in h^2^/L, but have similar L. Consequently, the joint distribution of z-scores and MAFs of their hits are markedly different (2 left panels), as are the marginal distributions of hit effect sizes, MAFs and z-scores (right). **(C)** After scaling by their respective 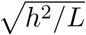, and imposing the more stringent scaled significance threshold (corresponding to FEV1) for both traits, the joint distribution of z-scores and MAFs of their hits (2 left panels) and the corresponding marginal distributions (3 right panels) are highly similar.

In contrast, height and FEV1 have similar mutational target sizes, but *h*^2^*/L* for height is 4.5-fold higher than for FEV1 (Figure 7B). As expected, this results in greater power to detect variants associated with height than with FEV1 (40% vs. 12% of *h*^2^ explained by significant hits). Consequently, genome-wide significant hits for height have larger effect sizes and larger *z*-scores than for FEV1. We also see slightly fewer low-MAF hits for FEV1 (p = 0.01, KS test), due to the lower power compared to height.

We predicted that after rescaling the z-scores (and effect sizes) for these traits by 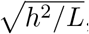, their architectures should become nearly identical. On this scale, the significance cutoff is 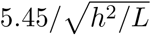, which is higher for FEV1 than for height (illustrated by the light and dark blue regions in Figure 6C). We therefore compared the architecture of genome-wide significant hits for both traits using the higher threshold in the scaled units. After doing so, we only have 131 hits for height, but it is apparent that the summary properties for both traits are indeed highly similar (Figure 7C).

We repeated this scaling procedure for the 50 other traits in our dataset whose scaled threshold is below that of FEV1. After doing so, the distributions for all hits are highly similar to the scaled distributions for height and FEV1 (dotted line, Figure 7C). We repeated similar analyses for all traits, and found these scaling laws approximate the architecture for all 95 traits (SI Section 8).

## Discussion

What determines the genetic architecture of complex traits? Does genetic variation in a given trait primarily reflect the idiosyncrasies of its biology–or alternatively, does it reflect processes that are shared among different traits?

Here we describe a principled approach to tackle these questions. Our point of departure is an evolutionary model of genetic architecture based on empirically motivated and interpretable biological assumptions. Given extensive evidence that many quantitative (continuous) traits are subject to stabilizing selection [23], and that genetic variation affecting one trait often affects many others [5,20,21], we model selection on alleles that arises from stabilizing selection in multi-dimensional trait space. Otherwise, we assume the standard population genetic model incorporating the effects of mutation, selection, genetic drift and demographic history. This model gives rise to a family of genetic architectures, where the architecture of a given trait is determined by the distribution of selection coefficients at trait-affecting sites as well as two scaling parameters: the mutational target size and the heritability per site.

We perform inference allowing all these parameters to vary among traits. We find that the model provides a good fit to the joint distribution of allele frequencies and effect sizes at genome-wide significant hits for the 95 quantitative traits in our dataset. Intriguingly, we also find that the distribution of selection coefficients is similar across all these traits, suggesting that their genetic architecture can be approximated by a Single Shared Distribution (SSD) of selection coefficients and two additional scaling parameters per trait. We then estimate the SSD using the data from all traits jointly, show that it fits the architecture of individual traits well, and validate our estimate of the SSD by showing that it accurately predicts the distribution of allele ages at GWAS hits for all traits.

The fit of the SSD model implies that, aside from the two scaling parameters, the genetic architecture is highly similar among all 95 traits. Cross-sections of the estimated shared architecture are visualized in Figure 6. Indeed, as predicted, after we rescale the effect sizes by the estimated heritability per site, we find that the joint distributions of effect sizes and allele frequencies for GWAS hits are remarkably similar among traits. Meanwhile the number of hits for a given trait is proportional to estimated target size (Figure 7).

These findings delineate the attributes of genetic architecture that are shared among traits and those that are trait-specific. In doing so, they raise new questions and insights about complex traits, as follows.

### Why is the distribution of selection coefficients similar across highly polygenic traits?

The similarity in genetic architecture among traits arises from the similarity in the distribution of selection coefficients of variants affecting them. Previous work has hinted at these similarities: for example, work using the *α*-model reported broadly similar relationships between MAF and effect size among a variety of traits [25, 26, 36], though it should be noted that there is no straightforward interpretation of *α* in terms of selection coefficients [36]. Why should the distribution of selection coefficents be similar across highly polygenic traits?

In the extreme, this similarity might be viewed as a consequence of high polygenicity and the finite amount of functional genetic variation. If all functional variants affected all traits, the distribution of selection coefficients would necessarily be shared among traits. This logic may well explain similarities among traits whose mutational target sizes encompass much of the functional portion of the genome; for example, we estimate the target size for BMI at ∼5% of the genome compared to ∼8% estimated to be functional [53–55]. The same logic may be especially relevant for traits that are mediated through the same tissues. However, this logic cannot explain the similarity among traits whose target sizes are substantially smaller and are primarily mediated through different tissues [56]. The biomarkers in our dataset, for example, have target sizes that are more than an order of magnitude smaller than BMI (e.g., calcium level with a target size of ∼0.1% of the genome), and are mediated through distinct cell types or tissues [9]. If traits are affected by different tissues, the variants affecting them should be different, so why should genetic variation in these traits be subject to similar selection?

One possibility is that the similarity in selection acting on variation affecting different traits reflects similarity in the biological systems in which variation arises, notably in gene-regulatory networks. Heritable variation in complex traits is spread across most of the genome, and is enriched in regulatory regions near most genes that are expressed in the tissues that affect these traits [14]. The lead variants typically explain a tiny fraction of the heritability [3], and the most relevant biological pathways are usually only modestly enriched for heritability [9, 14]. In other words, most heritable variation is mediated through the regulation of genes and pathways that are not closely connected to the trait’s biology [9, 14, 57]. Perhaps, the essential logic of gene regulatory networks and their evolution are sufficiently similar across tissues to drive similar distributions of selection coefficients, even if the specific pathways, genes, enhancers, and variants differ.

### Limitations and future analyses

We emphasize again that the SSD model is an approximation. There are indeed modest differences in the distributions of selection coefficients among traits as hinted at by the slightly better fits of the TSDs compared to the SSD. Additionally, our inference has limited power at both ends of the distribution of selection effects; hence we cannot exclude the possibility that there are larger differences among traits in these regions of the parameter space. Specifically, our focus on variants with MAF*>*1% limits our ability to quantify the contribution of very strongly selected, rare alleles, and focusing on genome-wide significant hits limits our analysis of weakly selected variants. Moreover, we only included traits with more than 100 genome-wide significant hits, thus limiting the scope of our results to highly polygenic traits. All these limitations may be addressed in the future. For example, other approaches [28], or whole genome sequencing [58] would enable greater power for strongly selected variants. Meanwhile, methodologies that integrate over the full distribution of causal variants accounting for LD [11, 19, 36] may allow us to relax the reliance on genome-wide significant hits, thus increasing the power to identify weakly selected variants, more generally increasing the precision of our inference for both the TSD and SSD models, and allowing for the analysis of less polygenic traits.

Future increases in sensitivity and applications of our inference to different kinds of traits may also warrant extensions of our evolutionary model. Notably, our current model is restricted to quantitative traits and is not immediately applicable to binary traits including diseases. Disease risk is often modeled in terms of an underlying quantitative trait referred to as liability, and for some diseases it seems plausible that the liability is subject to stabilizing selection (e.g., obesity with BMI as the underlying liability). We speculate that with appropriate adjustment of the model in order to fit data from case-control GWASs, we may find that such diseases share the same architecture as the quantitative traits that we studied here. The liability associated with other complex diseases, however, may plausibly be affected by directional selection to reduce disease risk, which is not captured in our current model. More generally, we assume that selection on variants arises solely from stabilizing selection in a high-dimensional trait space. While stabilizing selection may well be ubiquitous, one can imagine that some complex traits are subject to other modes of selection, such as the aforementioned directional selection to reduce disease risk. Another simplifying assumption of our model is that all the genetic variation in a given trait is affected by the same degree of pleiotropic selection (this degree was reflected in the dimension of the trait space; see, e.g., Equation 3 and Simons et al. 2018). Future extensions of the model may incorporate variation in the degree of pleiotropic selection, where this variation may also differ among complex traits.

### Why do trait-specific scaling parameters vary?

Future refinements notwithstanding, our evolutionary model with a shared distribution of selection effects fits the data from all 95 traits in our dataset remarkably well, indicating, as we have also confirmed, that differences in genetic architecture among highly polygenic traits are largely determined by the two trait-specific scaling factors: the mutational target size and heritability per site.

The mutational target size varies over 2 orders of magnitude among our 95 traits. While the estimates are novel, the variation among them is hardly surprising given the vast differences in the biology of these traits. These traits vary in being affected by few to many tissues, by the number and properties of core genes and pathways in these tissues [57], and by the number of independent biological processes that contribute to trait variation [59]. Moreover, the pathways associated with these traits plausibly differ in how buffered they are against genetic (and environmental) variation and in their modularity, plausibly reflective of the kinds of traits and of selection pressures over much longer evolutionary timescales than the turnover time of heritable variation [60, 61]. Our estimates of heritability are less variable, but still spanning over an order of magnitude, and vary among different kinds of traits (Table S1). Both of these observations have been known for almost a century, and yet the question about their causes remains largely open [23, 37, 38]. Our results indicate that other differences in architecture among traits are dwarfed by the variation caused by these two scaling parameters.

### Outlook

Taking a step back, our results highlight that evolutionary thinking is essential to understanding of the findings emerging from human GWASs, and more generally, heritable variation in complex traits. This insight is consistent with long-standing thinking in the field, given that heritable variation in complex traits reflects the outcome of evolutionary processes of mutation, natural selection, genetic drift and demographic history [23, 38]. Specifically, the signal measured in GWASs reflects causal variants’ contribution to heritability, which depends on their effect on the trait under consideration, but also on their minor allele frequency, where the relationship between the two is mediated by natural selection. What is surprising, at least to us, is how far an analysis based on evolutionary modeling can go, in this case, showing that the genetic architectures of highly polygenic quantitative traits are largely shared. This finding carries many implications about human GWASs and their applications, some of which we plan to explore elsewhere. Alongside other evidence, it also hints at underlying biology that largely remains to be discovered, plausibly relating to properties of gene regulatory networks. We think that a combination of evolutionary reasoning alongside a systems approach to gene regulation would move us closer to answering the questions that have existed since the beginning of the field of genetics, about the mapping from genetic variation to phenotypes.

## Acknowledgments

We thank Arbel Harpak, Luke O’Connor, Roshni Patel, Molly Przeworski, Jeff Spence, the Graham Coop lab, Pritchard lab, and Sella lab for conversations and comments. This work was funded by the following NIH grants: R01 HG008140, R01 HG011432, R01 GM115889, and F32 HG011202.

