## Supplementary Information for "Simple scaling laws control the genetic architectures of human complex traits"

October 4, 2022

### 1 The Model

We will present here a description of the pleiotropic stabilizing selection model used in the paper. It is an extension, and to a certain degree reparameterization, of the model presented in Simons et al 2018 [**?**]. Like the 2018 model, we focus on one focal trait under stabilizing selection and assume that variants may have pleiotropic effects on other, non-measured traits that also experience stabilizing selection.

We describe how the selection on a site depends on its effect on all traits. This allows us to arrive at expressions for the distribution of allele frequencies and effect sizes at trait-affecting sites, conditional on their selection coefficients. We then consider the existence of $L$ such sites, with $L$ being the mutational target size for the trait in question, and we denote the distribution of selection coefficient at such sites as $f(s)$. We show that, if effect sizes are measured in units of the phenotypic standard deviation, the last parameter needed to fully characterize the architecture is the heritability over the target size, i.e. the mean contribution to heritability per site.

Thus, our model describes the number of segregating sites affecting the trait, their frequencies and their effect sizes using $L$ and $f(s)$ and $h^2/L$ - the number of trait-affecting sites, the distribution of their selection coefficients and each site's mean contribution to heritability.

#### 1.1 Selection on Traits

We want to look at traits under stabilizing selection, that is when extreme trait values have lowered fitness compared to some medium, optimal trait value. We denote the trait value as $Y$ and we parametrize fitness as

$$W(Y) = \exp\left(-Y^2/2V_S)\right)$$

where $V_S$ is the width of the fitness function around the trait optimum, which we denote as the value $Y = 0$. Since under stabilizing selection models, phenotypes are tightly distributed around the optimum then this functional form can approximate any smooth and symmetric fitness function around the optimum.

The phenotype itself is modeled as an additive sum of genetic and environmental effects. The phenotype of an individual is a sum over the genetic contribution of genetic variants at many trait-affecting sites plus an environmental effect:

$$Y = \sum_i b_i \cdot g_i + e$$

where $b_i$ is the effect size of the derived allele at site $i$, $g_i = 0, 1, 2$ is the genoype at site $i$ and $e \sim N(0, V_E)$ is the environmental effect.

There may be, and most probably is, more than one trait under stabilizing selection. We assume that stabilizing selection acts on $n$ such traits, which we denote as $Y_1$,$Y_2$...$Y_n$, and that the fitness is multiplicative, i.e.

$$W(Y_1, Y_2...Y_n) = \exp(-Y_1^2/2V_S) \cdot \exp(-Y_2^2/2V_S) \cdots \exp(-Y_n^2/2V_S).$$

This equation can take a more succinct form if we think of the phenotype as a point in an n-dimensional trait space, i.e. an individual's phenotype is represented as the vector $\vec{Y} = (Y_1, Y_2, \ldots Y_n)$. We can now write

$$W(\vec{Y}) = \exp\left(-\|\vec{Y}\|^2/2V_S\right)$$

with $\vec{Y} = (Y_1, Y_2, \ldots Y_n)$ being a representation of the phenotype (i.e., the phenotypic value at all n traits) as a vector in an n-dimensional Euclidean trait space. As we discuss in Simons 2018, the fitness function around a peak can always be brought to this form using a linear transformation of the traits. We assume all variants affect the same number of traits and that all traits are identical, see Simons 2018 for discussion of the robustness of our results to violations of these assumptions.

Phenotypes can now be modeled as a vector additive sum of genetic and environmental effects. The phenotype of an individual is a sum over the genetic contribution of genetic variants at many trait-affecting sites plus an environmental effect:

$$\vec{Y} = \sum_i \vec{b}_i \cdot g_i + \vec{e}$$

where $\vec{b}_i$ is the effect size (in trait space) of the derived allele at site $i$ – meaning that the projection of $\vec{b}_i$ on any given dimension is the effect size of the derived allele at site $i$ on the trait represented by that dimension. $g_i = 0, 1, 2$ is still the genotype at site $i$ and $\vec{e} \sim N(0, V_E \cdot I)$, with $I$ being the $n$-dimensional identity matrix.

Since traits are sums of the small additive contributions of many alleles they will be approximately normally distributed in the population. As discussed in Simons 2018, we can assume that trait mean is at the optimum, which is denoted as $\vec{0}$, since stabilizing selection keeps it is very close to the optimum. We can therefore denote $\vec{Y} \sim N(0, V_P \cdot I)$.

#### 1.2 Selection on Variants

How does selection on traits affect variant allele frequencies?

In order to answer this question, we will calculate the expected change in allele frequency per generation caused by stabilizing selection on traits. We do this by averaging over the possible genetic background for the three possible genotypes at a site. Let's look at single site. We can now separate the phenotype into three contributions: site $i$, the genetic background (all other sites), and the environmental contribution:

$$\vec{Y} = \vec{b}_i \cdot g_i + \sum_{j \neq i} \vec{b}_j \cdot g_j + \vec{e} = \vec{b}_i \cdot g_i + \vec{Y}_{\text{not i}}$$

where $\vec{Y}_{\text{not i}}$ captures all effects other than site $i$.

What is the distribution of $\vec{Y}_{\text{not i}}$? The contribution from site i has mean $\vec{b}_i \cdot 2q_i$ and variance $\text{diag}(\vec{b}_i)^2 \cdot 2q(1-q)$. Together, the contribution from site $i$ and $\vec{Y}_{\text{not i}}$ have mean $\vec{0}$ and variance $V_P \cdot I$ and therefore we can approximate

$$\vec{Y}_{\text{not i}} \sim N(-\vec{b}_i \cdot 2q_i, V_P \cdot I)$$

where we assume the contribution to variance from site $i$, $\text{diag}(\vec{b}_i)^2 \cdot 2q(1-q)$, is small compared to $V_P$.

We can now calculate the mean fitness of each one of the three possible genotypes at site i. Let's look at individuals who are homozygous for the ancestral allele. Their phenotypes are just

$$\vec{Y}_{00} = \vec{Y}_{\text{not i}}$$

and so their mean fitness is

$$W_{00} = E\left[W(\vec{Y}_{\text{not i}})\right]$$

with $W(\vec{Y}) = \exp(-|\vec{Y}|^2/2V_S)$, as before, and the expectation taken over the distribution of $\vec{Y}_{\text{not i}}$. Similarly, for the heterozygote $\vec{Y}_{01} = \vec{Y}_{\text{not i}} + \vec{b}_i$ and therefore

$$W_{01} = E\left[W(\vec{Y}_{\text{not i}} + \vec{b}_i)\right]$$

and similarly

$$W_{01} = E\left[W(\vec{Y}_{\text{not i}} + 2\vec{b}_i)\right]$$

and the first moment of change in allele frequency is

$$E[\Delta q] = -pq\frac{p(W_{00} - W_{01}) + q(W_{01} - W11)}{\bar{W}}$$

with $\bar{W} = p^2 W_{00} + 2pqW_{01} + q^2 W_{11}$.

The expression for $E[\Delta q]$ greatly simplifies when $|\vec{b}_i|^2 \ll V_S$ and $V_P \ll V_S$, i.e. when the reduction in mean log fitness due to site $i$ and the overall phenotypic variation is much smaller than 1. In this case the relative fitnesses are

$$\frac{W_{00}}{\bar{W}} \approx 1 - \frac{4q^2 \cdot |\vec{b}_i|^2}{2V_S}, \; \frac{W_{01}}{\bar{W}} \approx 1 - \frac{(p-q)^2 \cdot |\vec{b}_i|^2}{2V_S}, \; \frac{W_{11}}{\bar{W}} \approx 1 - \frac{4p^2 \cdot |\vec{b}_i|^2}{2V_S},$$

and

$$E[\Delta q] \approx -\frac{|\vec{b}_i|^2}{V_S} \cdot pq(\tfrac{1}{2} - q) = -s \cdot pq(\tfrac{1}{2} - q)$$

and we see that $s = \frac{|\vec{b}_i|^2}{V_S}$ serves as the selection coefficient acting on a variant.

Note that $E[\Delta q]$ takes the classic form for underdominant selection even though the relative fitnesses of the three genotypes do not. What we observe is an effective underdominant selection that comes about through the frequency dependency of the relative fitnesses.

The above equation for $E[\Delta q]$, together with demography, determines the distribution of derived allele frequencies for variants with a given selection coefficient, which we denote as $P(q|s)$. This distribution can be estimated via forward simulations (as we do) or solving the relevant Kolmogorov equation. [1]

[1]Although the Kolmogorov equation can usually only be solved numerically, for a constant population size an

##### 1.3 Relationship between selection coefficients and effect sizes

Under our model, the strength of selection acting on a variant, i.e. its selection coefficient, is determined by the variant's phenotypic effect on all traits. However, for solving the model, it is convenient to express the distribution of effect size on a focal trait conditional on the selection coefficient, i.e. conditional on the overall effect on all traits. We obtain an expression for this distribution using purely geometric reasoning.

Variants with a given n-dimensional effect size correspond to a hypersphere in the $n$ dimensional trait space with radius $|\vec{b}|$, see fig. ??A. Of those, variants with a given effect size on our focal trait, $b_1$, correspond to a cross-section of that sphere. The ratio of the cross-section's area to the sphere's area is the density of variants with effect size $b_1$.

We will now calculate this ratio for variants with effect size between $b_1$ and $b_1 + \Delta b_1$, with $\Delta b_1$ being arbitrarily small. The area of that sphere is proportional to its radius to the $(n-1)$th power, i.e. to $|\vec{b}|^{n-1}$. The area of the cross-section is proportional to its radius, which is $\sqrt{|\vec{b}|^2 - b_1^2}$, to the power of $n-2$ times $\Delta b_1 \cdot \frac{|\vec{b}|}{\sqrt{|\vec{b}|^2 - b_1^2}}$ , the arc length corresponding to $\Delta b_1$, see Fig. ??B.

Therefore, the ratio of the areas is

$$\frac{A_{\text{cross section}}}{A_{\text{sphere}}} \propto \frac{\left(\sqrt{|\vec{b}|^2 - b_1^2}\right)^{n-2} \cdot \Delta b_1 \cdot \frac{|\vec{b}|}{\sqrt{|\vec{b}|^2 - b_1^2}}}{|\vec{b}|^{n-1}} = \left(\sqrt{1 - \frac{b_1^2}{|\vec{b}|^2}}\right)^{\frac{n-3}{2}} \cdot \frac{\Delta b_1}{|\vec{b}|}$$

The density of $b_1$ conditional on $|\vec{b}|$ will then be

$$f(b_1) \propto \left(\sqrt{1 - \frac{b_1^2}{|\vec{b}|^2}}\right)^{\frac{n-3}{2}} \cdot \frac{1}{|\vec{b}|}.$$

This somewhat daunting expression greatly simplifies when $n \gg 1$. In this limit, we know that the squared effect on the focal trait is of the order of $|\vec{b}|^2/n$ making $b_1^2/|\vec{b}|^2 \sim O(1/n)$. In addition, since $n \gg 1$ then $(n-3)/2 \approx n/2$. Taking these simplifications together we get that

$$f(b_1) \propto \left(\sqrt{1 - \frac{b_1^2}{|\vec{b}|^2}}\right)^{\frac{n-3}{2}} \cdot \frac{1}{|\vec{b}|} \approx \left(\sqrt{1 - \frac{b_1^2}{|\vec{b}|^2}}\right)^{\frac{n}{2}} \cdot \frac{1}{|\vec{b}|} \approx \exp\left(-\frac{n}{2} \cdot \frac{b_1^2}{|\vec{b}|^2}\right) \cdot \frac{1}{|\vec{b}|} = \frac{1}{\sqrt{|\vec{b}|^2}} \cdot \exp\left(-\frac{1}{2} \cdot \frac{b_1^2}{|\vec{b}|^2/n}\right).$$

which we immediately recognize as a Gaussian function meaning that

$$b_1 \sim N\left(0, \frac{|\vec{b}|^2}{n}\right).$$

Since effect sizes on a focal trait are normally distributed conditional on the $n$-dimensional effect, they are also normally distributed conditional on the selection coefficient (since $s = |\vec{b}|^2/V_S$):

$$b_1 \sim N\left(0, \frac{V_S}{n} \cdot s\right).$$

analytical solution exists, see Simons 2018. For a large population size, it can be shown that this analytical solution for the minor allele frequency takes the simple form

$$P(x|s) = 2N_e\mu \cdot \frac{2\exp(-2N_e s \cdot x(1-x))}{x(1-x)}$$

for $x > 1/2N_e$, with $x$ being the minor allele frequency. Though we don't use it in this work we thought it could be a useful result to mention.

Because this result relies on the geometry of high dimensional spaces it holds even when we relax the assumption of complete isotropy, albeit with an effective dimension replacing n in the above equation. See Simons (2018) for more details. Therefore, when looking at a given trait, we can drop the subscript of $b_1$ and arrive at

$$P(b|s) = \frac{1}{\sqrt{2\pi V_S/n}} \cdot \exp\left(-\frac{1}{2} \cdot \frac{b_1^2}{V_S/n}\right).$$

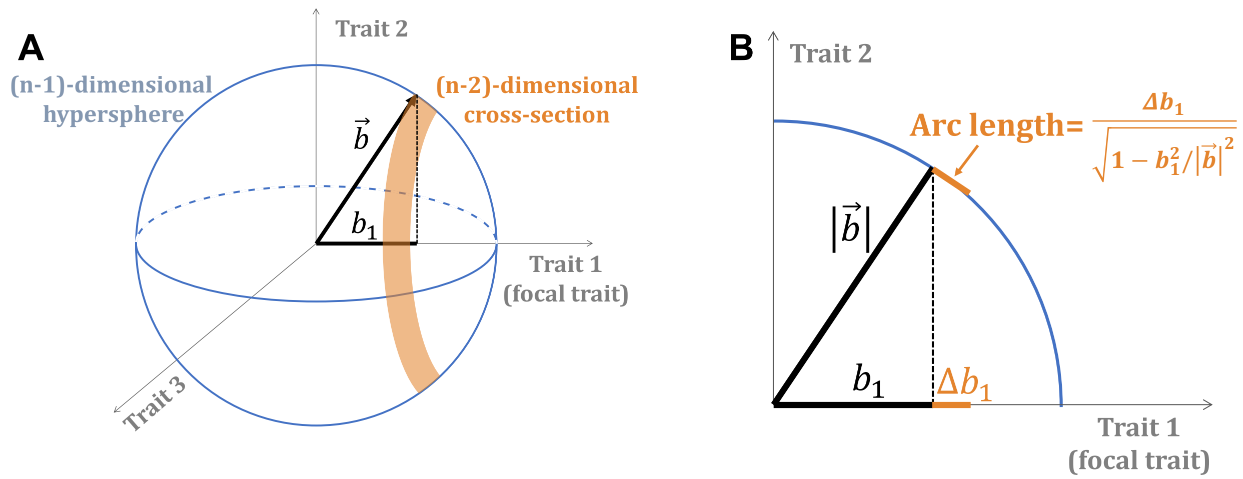

Figure S1: **Geometric relationship between the $n$-dimensional effect and its projection on a single trait.** ***(A)*** *The possible values of $\vec{b}$ for a given $|\vec{b}|$ form an $n-1$-dimensional hypersphere in the $n$ dimensional trait space. The values of $\vec{b}$ corresponding to a given effect size on the focal trait, $b_1$, form a cross-section of the hypersphere.* ***(B)*** *The geometry of calculating the arc length of the cross section.*

#### 1.4 Genome-wide architecture

We can now combine the above results to calculate the distribution of derived allele frequency, $q$, and effect size, $b$, for $L$ sites affecting the trait (i.e., $L$ is the mutational target size for the trait). The density of sites with a given $q$ & $b$ for a given selection coefficient $s$ is given by

$$P(q, b|s) = P(q|s) \cdot P(b|s)$$

with $P(q|s)$ and $P(b|s)$ as given above. Note that, conditional on the selection coefficient, the frequency and effect size are independent.

The overall distribution of $q$ & $b$ at all $L$ sites is given averaging over the distribution of selection coefficients $f(s)$

$$P(q, b) = \int_s P(q, b|s) \cdot f(s) \cdot ds$$

Averages over the distribution of frequencies and effect sizes can then be calculated as

$$E[g(q, b)] = \int_q \int_b g(q, b) \cdot P(q, b) \cdot dbdq$$

with $g$ being any function of $q$ & $b$. Expectation of sums over all sites can be calculated by

$$E\left[\sum_i g(q_i, b_i)\right] = L \cdot E[g(q, b)].$$

To be concrete we will give a few useful examples:

- The probability of a variant being above a given frequency threshold $q^*$ is
  $P(q > q^*) = E\left[\mathbb{1}_{q>q^*}\right] = \int_q \int_b \mathbb{1}_{q>q^*} \cdot P(q,b) \cdot dbdq = \int_{q>q^*} \int_b P(q,b) \cdot dbdq$
  and therefore the number of variants above that threshold is
  $\#_{q>q^*} = L \cdot P(q > q^*)$.
- The mean effect on the phenotype of variation at site is
  $E[2q \cdot b] = \int_q \int_b 2q \cdot b \cdot P(q,b) \cdot dbdq$
  and the overall mean genetic contribution to the phenotype is
  $E[Y] = L \cdot E[2q \cdot b]$.
  Without mutational bias or directional selection both of these expected values are equal to zero.
- The mean contribution to phenotypic variance from a site is
  $E[2q(1-q) \cdot b^2] = \int_q \int_b 2q(1-q) \cdot b^2 \cdot P(q,b) \cdot dbdq$
  and therefore the genetic contribution to variance is
  $V_G = L \cdot E[2q(1-q) \cdot b^2]$.

##### 1.5 Reparameterization in terms of $h^2/L$

) When looking at single sites, we have expressed the relation between selection and effect size in terms of the width of the fitness function around the optimum, $V_S$, and the degree of pleiotropy, i.e. number of traits $n$. Looking at all sites, we can express this relation using the heritability $h^2$ and the target size $L$, allowing us to better interpret our results.

Under our model, the contribution to genetic variance from all sites is

$$V_G = L \cdot E[2q(1-q) \cdot b^2]$$

and when measuring effect size in units of the phenotypic standard deviation this equation scales to

$$h^2 = V_G/V_P = L \cdot E[2q(1-q) \cdot \beta^2]$$

with

$$\beta = b/\sqrt{V_P}.$$

Therefore,

$$\frac{h^2}{L} = E[2q(1-q) \cdot \beta^2]$$

and we see that the heritability over the target size is the expected contribution to heritability from a single site, i.e. the heritability per site. Conditional on the selection coefficient, effect sizes and allele frequencies are independently distributed. We can take the expectation over the distribution of effect sizes conditional on the selection coefficient – since $E\left[b^2|s\right] = (V_S/n) \cdot s$ then $E\left[b^2|s\right] = (V_S/n \cdot V_P) \cdot s$. Therefore,

$$\frac{h^2}{L} = \frac{V_S}{n \cdot V_P} E[2q(1-q) \cdot s] = \frac{E[2q(1-q) \cdot s]}{n \cdot V_P/V_S}.$$

We have now arrived at a second interpretation of the heritability per site. For small selection coefficients, the numerator on the right hand side of the above equation is the mean reduction in mean log fitness due to a single site. The denominator the reduction in mean log fitness due to the overall variation in phenotype (in all n traits). So the heritability per site is both the mean relative contribution of single site to genetic variance and the mean relative contribution of a single site to the reduction in log fitness.

Under mutation-selection balance $E[2q(1-q)\cdot s] = 4\mu$, with $\mu$ being the mutation rate. As you can see in figure **??**A, this relation approximately holds for single selection coefficients as long as selection is comparable with genetic drift under a European-like demographic history (see in depth discussion of the effects of demography in [**?**]). Therefore, the relationship $E[2q(1-q)\cdot s] \approx 4\mu$ holds in general (see **??**B), albeit with a small dependency on the distribution of selection coefficients.

We can therefore write

$$\frac{V_S}{n \cdot V_P} = \frac{k}{4\mu} \cdot \frac{h^2}{L}$$

with $k \approx 1$ and therefore

$$\beta \sim N\left(0, \frac{k}{4\mu} \cdot \frac{h^2}{L} \cdot s\right)$$

and we see that the heritability per site sets the scale of effect sizes in units of the phenotypic standard deviation.

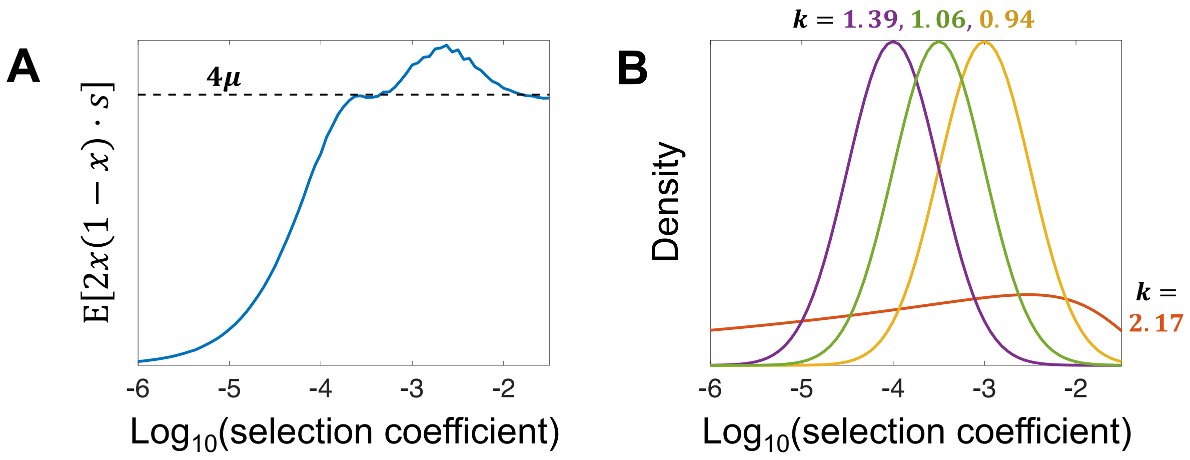

Figure S2: **Mutation-selection balance approximation.** **(A)** *For a single selection coefficient, the approximation $E[2q(1-q)\cdot s] \approx 4\mu$ holds for $s \geq 10^{-4}$ even with non-esquilibrium demography. The scale of $E[2q(1-q)\cdot s]$ is set by $4\mu$ even for $s < 10^{-4}$.* ***(B)*** *As a result, we can define $E[2q(1-q)\cdot s] = 4\mu/k$ for a distribution of selection coefficient, and we see that for all distributions $k$ is of the order of 1.*

#### 1.6 Summary of model

We have described a model with three inputs: the number of trait-affecting sites, or mutational target size, denoted as $L$; The distribution of selection coefficients at trait-affecting sites, denoted as $f(s)$; and the mean contribution to heritability of a trait-affecting site, denoted as $h^2/L$. New mutations arise at the $L$ sites, thus sampling selection coefficients from $f(s)$.

The distribution of derived allele frequencies at a site is determined by the selection coefficient and the population's demography and we denote it as $P(q|s)$. The distribution of effect sizes (in units of the phenotypic standard deviation), which we denote as $P(\beta|s)$, is a normal distribution with mean zero and variance approximately equal to $1/4\mu \cdot h^2/L \cdot s$. The distribution of any summary of frequency and effect sizes follows from these distributions.

### 2 The Likelihood

We will present here an overview of our method to infer the three components of our model - $L$, $f(s)$ and $h^2/L$ - from the co-distribution of minor allele frequencies and z-scores of GWAS hits. To do this, we first recast our model from derived allele frequencies and effect sizes to minor allele frequencies and z-scores, which allows us to condition on variants being GWAS hits. We then write a composite log-likelihood for $L$, $f(s)$ and $h^2/L$ given the minor allele frequencies and z-scores of GWAS hits. Lastly, we tweak this likelihood to account for varying imputation qualities among SNPs.

#### 2.1 Minor allele frequencies

In order to simplify our equations and avoid having to account for errors in identifying the derived allele frequency of the causal variant tagged by a GWAS hit, we fold the frequency spectrum, i.e. work exclusively with minor allele frequencies (MAFs) in our likelihood. The minor allele frequency is defined as

$$x = min(q, 1-q)$$

and its distribution is

$$P(x|s) = P(q|s) + P(1-q|s).$$

The MAF can take values between 0 and 0.5. Due to the decline in imputation quality at low MAFs, we will restrict ourselves to sites with $x > 0.01 = 1\%$. At frequencies above 1% and with study sizes above 100k, the estimation error for $x$ is very small ($\Delta x/x \leq 0.003$). Therefore, we ignore the estimation error and assume the estimate MAF is the true MAF for each site.

#### 2.2 The estimated effect size

With study sizes in the hundreds of thousands estimation error in MAFs is insignificant, but errors in estimating effect sizes are not. With study size $M \gg 1$, the estimated effect size, $\hat{\beta}$, as a function of the true effect size, $\beta$, at an allele of minor allele frequency $x$ is

$$\hat{\beta}|\beta, x \sim N\left(\beta, \frac{1/M}{2x(1-x)}\right)$$

with effect sizes measured in units of the phenotypic standard deviation ($V_P$).

For a given selection coefficient, not only are the estimation errors approximately normally distributed but so is the true effect size

$$\beta \sim N\left(0, \frac{k}{4\mu} \cdot \frac{h^2}{L} \cdot s\right).$$

Therefore, estimated effect sizes at sites with selection coefficient $s$ and MAF $x$ are normally distributed as

$$\hat{\beta}|s,x \sim N\left(0, \frac{k}{4\mu} \cdot \frac{h^2}{L} \cdot s + \frac{1/M}{2x(1-x)}\right)$$

where the first term corresponds to the true effect and the second to the estimation error.

#### 2.3 z-score

Since genome-wide significance is defined by a cutoff on the z-scores of variants, it is convenient to work with the z-scores instead of the estimated effect sizes, i.e. normalize the estimated effect sizes such that the error term is identically equal to 1. By this definition,

$$z = \sqrt{M \cdot 2x(1-x)} \cdot \hat{\beta}$$

and therefore

$$z|s,x \sim N\left(0, \frac{k}{4\mu} \cdot \frac{M \cdot h^2}{L} \cdot 2x(1-x) \cdot s + 1\right).$$

Note that, unlike the effect size, the z-score depends on both the selection coefficient and the minor allele frequency.

We define a combined GWAS power parameter

$$C = \frac{k}{4\mu} \cdot \frac{M \cdot h^2}{L}$$

which captures the scale of true signal ($V_S/n$ or $\frac{k}{4\mu}V_G/L$) relative to the scale of estimation error ($V_P/M$). With this definition,

$$z|s,x \sim N\left(0, C \cdot 2x(1-x) \cdot s + 1\right).$$

It is this power parameter $C$, that we will directly infer in our inference. We denote the density function of the z-score as

$$P(z|C,s,x)$$

which is a normal distribution with mean zero and variance $C \cdot 2x(1-x) \cdot s + 1$.

#### 2.4 Varying imputation quality

Ideally, the genotype at all sites would be perfectly estimated for all individuals. In practice, many sites are imputed and not directly genotyped. Even sites that are directly genotyped might be missing for some individuals. The result is that the study size, $M$, in the equations above varies between sites. We define the effective study size for site $i$ as

$$M_i = (\Delta\beta_i)^2 \cdot 2x(1-x)$$

with $\Delta\beta_i$ being the reported standard error in effect size (usually estimated by resampling). We find the median study size over all SNPs with x¿1%, including non-significant SNPs, as $M_{med} = \text{Median}(\{M_i\})$. $M_{med}$ is nearly identical to the reported study size for each trait.

We define for each site a relative study size

$$m_i = \frac{M_i}{M_{med}}.$$

This relative study size is trait-independent and is proportional to (and essentially determined by) the info score that measures imputation quality. For included SNPs, $m$ ranges from 0.8 to just over 1. $m$ has a denoted dependency on the minor allele frequency and we estimate, for all SNPs, the distribution

$$P(m|x)$$

for different frequency bins (see bin definition in the next Section) using the standard errors for height effect sizes.

We now have

$$z|s,x \sim N\left(0, m \cdot C \cdot 2x(1-x) \cdot s + 1\right).$$

and the corresponding

$$P(z|m,C,s,x).$$

#### 2.5 The joint distribution of MAFs and z-scores

We can now write expressions for the distributions of MAFs, z-scores among trait-affecting variants. Conditional on the selection coefficient, and power C the distribution of MAFs and z-score, and effective study sizes is

$$P(x,z,m|C,s) = P(z|m,C,s,x) \cdot P(m|x) \cdot P(x|s)$$

and note that $x$ and $z$ are not independent, rather $z$ is dependent on $x$. We can integrate over the effective study sizes to arrive at the distribution of MAF and z-scores for each selection coefficient

$$P(x,z|C,s) = \int_m P(z|m,C,s,x) \cdot P(m|x) \cdot P(x|s) \cdot dm$$

and lastly we can integrate over the selection coefficients to arrive at the overall distribution of $x$ & $z$

$$P\left(x,z|C,f(s)\right) = \int_s P(x,z|C,s) \cdot f(s) ds.$$

We can take expectations over this distribution

$$E\left[g(x,z)|C,f(s)\right] = \int_x \int_z g(x,z) \cdot P\left(x,z|C,f(s)\right) \cdot dxdz.$$

and get expectation of quantities summed over all sites by

$$E\left[\sum_i g(x_i,z_i)|C,f(s)\right] = L \cdot E\left[g(x,z)|C,f(s)\right].$$

#### 2.6 Threshold for genome-wide significance

We limit our inference to use GWAS hits, i.e. genome-wide significant variants. While it is difficult to estimate the MAF and effect size of causal effect sizes, there exists methods to (approximately) infer a set of SNPs representing independent signals and each SNP should tag a single causal variant. Though such tagging SNPs are not themselves necessarily causal, they are in tight LD with the causal variants and therefore represent the causal variant's MAF and effect size.

A variant is considered genome-wide significant is its p-value under the null hypothesis of no effect size (null of $\beta = 0$) is below a threshold value, usually taken to be $5 \cdot 10^{-8}$. Since p-values are a monotonically decreasing function of the z-score squared, this threshold on the p-value translates to a threshold on the absolute value of the z-score. For a p-value threshold of $5 \cdot 10^{-8}$ this threshold is

$$|z| > z^* = \sqrt{2} \cdot \text{erfc}^{-1}\left(5 \cdot 10^{-8}\right) \approx 5.45$$

with $\text{erfc}^{-1}$ being the inverse complimentary error function (mercifully, we will not derive this formula here).

#### 2.7 The conditional co-distribution of MAFs and z-scores

We now want to write an expression for the distribution of $x$ and $z$ for GWAS hits with MAF above 1%.

First, let's calculate the probability of a variant being a hit at x¿1% with relative study size $m$

$$Pr\left(hit|m, C, f(s)\right) = E\left[\mathbb{1}_{|z|>z^*} \cdot \mathbb{1}_{x>1\%}|m, C, f(s)\right] = \int_x \int_z \mathbb{1}_{|z|>z^*} \cdot \mathbb{1}_{x>1\%} \cdot P\left(x, z, m|C, f(s)\right) \cdot dxdz.$$

and therefore the conditional distribution of $x$, $z$ and $m$ is simply

$$P(x, z, m|C, s) = \frac{P(x, z, m|C, s)}{Pr\left(hit|m, C, f(s)\right)}.$$

Note, that this distribution depends on the distribution of selection coefficients $f(s)$ and power parameter $C = M_med \frac{k}{4\mu} \frac{h^2}{L}$, but not on the target size $L$. The target sizes $L$ determines the expected number of GWAS hits via

$$E[\#_{hits}|C, f(s)] = L \cdot Pr(hit|C, f(s))$$

with

$$Pr\left(hit|C, f(s)\right) = E\left[\mathbb{1}_{|z|>z^*} \cdot \mathbb{1}_{x>1\%}|C, f(s)\right] = \int_x \int_z \mathbb{1}_{|z|>z^*} \cdot \mathbb{1}_{x>1\%} \cdot P\left(x, z|C, f(s)\right) \cdot dxdz.$$

#### 2.8 Estimating model parameters

We are finally ready to write down our likelihood and estimators for our model parameters.

Since the distribution of $x$ and $z$ depends on the distribution of selection coefficients $f(s)$ and power parameter $C = M_{med} \frac{k}{4\mu} \frac{h^2}{L}$, we can write a composite log-likelihood for these parameters as

$$LL\left(C, f(s)|\{x, z, m\}\right) = \sum_i \log\left(P\left(x_i, z_i, m_i|hit, C, f(s)\right)\right).$$

Our estimates $\hat{f}(s)$ & $\hat{C}$ are those that maximize this likelihood. Our estimate of $h^2/L$ is

$$\frac{\hat{h^2}}{L} = E[s \cdot 2x(1-x)|\hat{C}, \hat{f}(s)] \frac{\hat{C}}{M_{med}} = \frac{4\mu}{k} \cdot \frac{\hat{C}}{M_{med}}.$$

The number of hits will be Poisson distribution with mean $L \cdot Pr(hit|C, f(s))$. Therefore, once we estimated $f(s)$ and $C$, the maximum likelihood estimator of $L$ is simply

$$\hat{L} = \frac{\#_{hits}}{Pr(hit|\hat{C}, \hat{f}(s))}$$

with $Pr(hit|\hat{C}, \hat{f}(s))$ estimated using $\hat{f}(s)$ & $\hat{C}$ .

##### 2.9 How do MAFs and z-scores translate to selection coefficients?

It is not obvious from the above expressions for the likelihood how do the MAF and z-scores translate to selection coefficients. However, with a bit of analysis it is easy to see that, for each variant, the MAF provides an upper bound on the selection coefficient while the z-score provides a lower bound.

Let's think first of a population with a constant population size: When selection is as strong as (or stronger than) genetic drift, the MAFs of variants are limited to MAFs below $1/2N_es$, see Fig. **??**A. Therefore, the likelihood of a variant at a given MAF $x$ coming for selection coefficient $s$, is flat for $s < 1/2N_ex$ and falls dramatically for larger selection coefficients, see Fig. **??**B. This same picture holds qualitatively for non-equilibrium demography too.

Under our model, z-scores squared are of the order of $C \cdot 2x(1-x) \cdot s$, i.e. larger for larger selection coefficients, see Fig. **??**C. This immediately suggests that the log-likelihood of $s$ for a given $z$ falls when $s$ becomes being larger than $z^2/C \cdot 2x(1-x)$, as we can indeed see in Fig. **??**D.

Taken together, the MAF provides an upper bound on the selection coefficients and the z-score provides a lower bound, Fig. **??**E. Though these bounds still leave a lot of uncertainty for each variant, taken together even 100 variants are enough to estimate the distribution of selection coefficients.

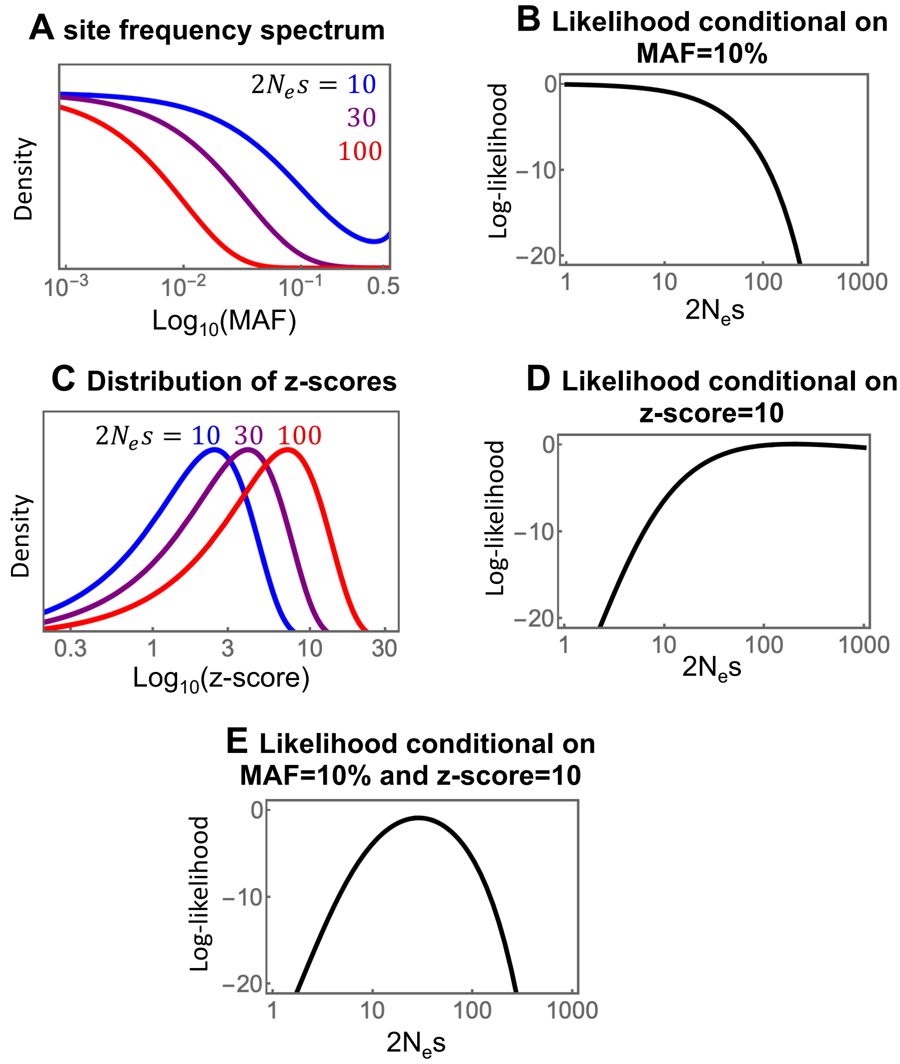

Figure S3: **MAF and z-score bound the selection coefficient.** Illustration of how MAF and z-score bound the selection coefficient in a simplified scenario of constant population size and perfect knowledge of causal variants. ***(A)*** *The probability density of MAFs for different strengths of selection (the SFS).* ***(B)*** *The resulting log-likelihood of selection coefficients given a MAF of 10%.* ***(C)*** *The distribution of z-scores for different strengths of selection.* ***(D)*** *The resulting log-likelihood of selection coefficients given a z-score of 10.* ***(E)*** *The log-likelihood of selection coefficients given MAF of 10%* ***and*** *a z-score of 10. (We took $N_e = 10,000$ and $C = 5,000$.)*

### 3 Maximizing the likelihood

While writing down expressions of the likelihood is rather straightforward, reliably estimating this likelihood with a reasonable runtime has proven to be quite a challenge.

We then discuss a few practical hurdles to inferring our model's parameters. First, we explain how we estimate the likelihood in practice: We parametrize $f(s)$ as a log-spline and discretize some of the underlying parameters. Second, we explain our three step approach to maximizing the likelihood and inferring the model's parameters: we find the best fitting $f(s)$ using simulated annealing. At each iteration, given $f(s)$, we perform a simple line search over $h^2/L$. Once $f(s)$ and

$h^2/L$ are estimated, we use those estimates and the number of GWAS hits to estimate $L$. Lastly, we discuss the necessary addition of a regularization penalty on $f(s)$ to the likelihood.

##### 3.1 Binning and gridding

In order to handle the calculations involved in estimating model parameters we treat $x$,$s$ and $C$ as discrete parameters. This allows us to formulate our equations as linear operations on probabilities. Since the z-score is normally distributed conditional on the other parameters, calculating the z-score's probability density is straightforward and we do not need to discretize it.

In order to bin the minor allele frequencies, we use 26 bins between 1% and 50%. The bin limits are:

$$\{10^{-2}, 10^{-1.9}, 10^{-1.8}, 10^{-1.7}, 10^{-1.6}, 10^{-1.5}, 10^{-1.4}, 10^{-1.3}, 10^{-1.2}, 10^{-1.1}, 0.1$$
$$, 0.125, 0.15, 0.175, 0.2, 0.225, 0.25, 0.275, 0.3, 0.325, 0.35, 0.375, 0.4, 0.425, 0.45, 0.475, 0.5\}$$

We look at selection coefficients on a dense grid on a log scale from $s = 10^{-8}$ to $s = 10^{-1}$ in steps of $10^{1/16}$ (113 selection coefficients). We also consider the power parameter $C$ on a dense grid on a log scale from $C = 10^3$ to $C = 10^8$ in steps of $10^{0.01}$ (501 $C$ values).

##### 3.2 Site frequency spectra

For each of the selection coefficients on our grid above $10^{-6}$ we ran our forward simulator (see details in Simons 2018 [**?**]) 120 million times, simulating 120Mbp of sites with that selection coefficient. We used a demographic model inferred using RELATE [**?**] on the British population(GBR) in the 1000 genomes dataset , see Fig. **??**. Since the distribution of common allele frequencies becomes insensitive to the selection coefficients when they are very small, we used the simulation for $10^{-6}$ when considering smaller selection coefficients (we consider such tiny selection coefficients in order to account for their possible effect on the distribution of effect sizes).

Thus, for each selection coefficient we estimate the proportion of variants with MAF above 1% and the distribution of their MAFs. This allows us to estimate, the probability of a variant being at each frequency bin, which we use as a proxy for $P(x|s)$.

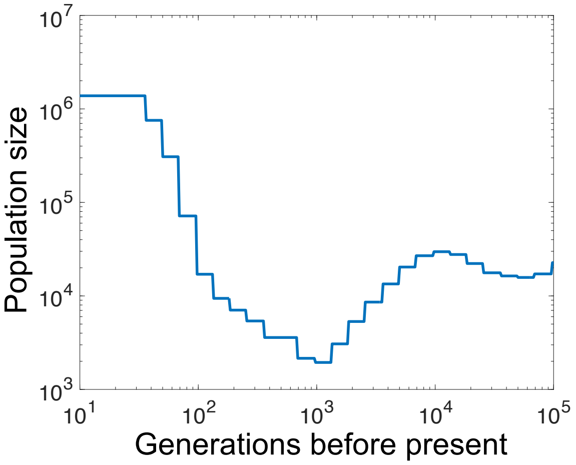

Figure S4: **Demographic model.** *The demographic model we use was inferred for the GBR 1000 genomes population using RELATE [**?**].*

##### 3.3 Precaculated probabilities

Estimating our model parameters begins by precalculating all necessary probabilities conditional on $s$ and $C$, for all $s$ and $C$ on our grid. For each GWAS hit $i$, we calculate the matrix

$$P^i_{C,s} \equiv P(x_i, z_i, m_i|C, s) = P(z_i|m_i, C, s, x_i) \cdot P(m_i|x_i) \cdot P(x_i|s).$$

This is a 501×113 matrix with three components:

- $P(z_i|m_i, C, s, x_i)$ - the z-score probability. This is just a normal pdf with variance $m_i \cdot C \cdot s \cdot 2x_i(1 - x_i) + 1$ estimated at $z_i$ for each value of $C$ and $s$.
- $P(m_i|x_i)$ - the probability of the relative study size. This is a single number that is independent of both $C$ and $s$. This number will just add a constant to the log likelihood and therefore would not affect our parameter estimates. We therefore replace it with 1.
- $P(x_i|s)$ - the SFS of variants,. This is the probability of seeing a variant with selection coefficient $s$ at MAF $x$, which we estimate using our forward simulations.

In addition, we precalculate the corresponding matrix of probabilities of variants being hits for all $C$ and $s$,

$$P^{hit}_{C,s} = E\left[\mathbb{1}_{|z|>z^*} \cdot \mathbb{1}_{x>1\%}|C, s\right] = \int_x \int_z \int_m \mathbb{1}_{|z|>z^*} \cdot \mathbb{1}_{x>1\%} \cdot P(z|m, C, s, x) \cdot P(m|x) \cdot P(x|s) \cdot dxdzdm.$$

We numerically calculate this 501×113 matrix monstrosity. Luckily, $P^{hit}_{C,s}$ is the same for all variants and traits so we only have to calculate it once.

##### 3.4 Calculating the likelihood via linear algebra

The advantage of precalculating all the probabilities, which takes an incredible amount of computation and memory, is that it is now extremely trivial to calculate the likelihood

$$LL\left(C, f(s)|x, z, m\right) = \sum_i \log\left(\frac{\sum_s P^i_{C,s} \cdot f_s}{\sum_s P^{hit}_{C,s} \cdot f_s}\right)$$

where we denote as $f_s$ the column vector of size 113 representing $f(s)$ at each selection coefficient on our grid ($\sum_s f_s = 1$).

##### 3.5 Estimating $C$ for a given $f(s)$

It is now trivial to estimate $C$ for a given $f(s)$. The likelihood conditional on $f(s)$ can be thought of as a row vector

$$LL(C) = \sum_i \log\left(\frac{\sum_s P^i_{C,s} \cdot f_s}{\sum_s P^{hit}_{C,s} \cdot f_s}\right)$$

and then

$$\hat{C} = \text{argmax}_C(LL(C))$$

and we denote the marginal log-likelihood at this estimate $\hat{C}$ as

$$LL\left(f(s)\right) = LL(\hat{C}) = \max_C LL\left(C, f(s)|x, z, m\right).$$

##### 3.6 Estimating $f(s)$ from the marginal likelihood

In principle, estimating $f(s)$ just involves maximizing $LL(f(s))$. In practice, in order to do that we need to somehow parametrize $f(s)$ with a parametrization that's both flexible and parsimonious. We do that via a log-spline, i.e. we parametrize the log of $f(s)$ as a spline, with $s$ on a log scale.

The log-spline is parametrized by its value at specific knots, we use 4 knots at $\{k_1, k_2, k_3, k_4\} = \{-6, -4.5, -3, -1.5\}$, and our parameters are the value of $\log f(s)$ at $s = \{10^{k_i}\}$. Our $f(s)$ for every value of $s$ is given by

$$\log f(s) = \text{spline}(\log_{10} s | \{k, \log f(10^k)\})$$

with spline here indicating a cubic spline from the 4 pairs $\{k, \log f(10^k)\}$ to $log_{10}s$. We estimate $f(s)$ at our 113 grid points to arrive at the vector $f_s$, which we normalize to 1. We then use simulated annealing to find the values of $\{\log f(10^k)\}$ that maximize $LL(f(s))$. These values define our estimate $\hat{f}(s)$. In Fig. **??**, we can see these inferred distributions for all 95 traits (with TSDs) and appreciate that they are very similar.

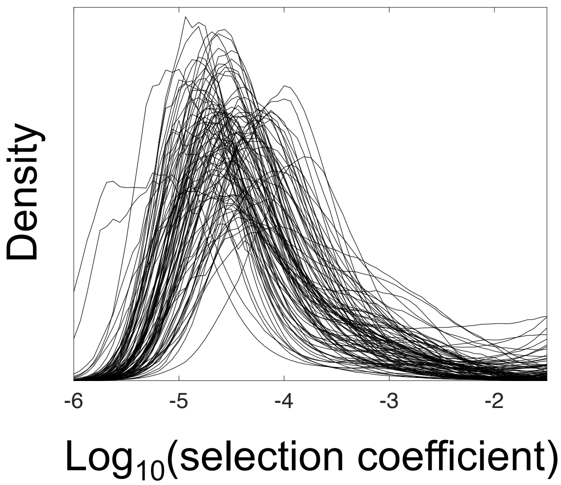

Figure S5: **The inferred distribution of selection coefficients for all 95 traits.** Presented here is the median of the distribution over 100 bootstraps.

##### 3.7 Maximizing the likelihood for a single shared distribution of $s$

So far, we have only considered one trait at a time, i.e. the trait-specific distributions (TSDs) model. Our setup is easily applicable to the single shared distribution model (SSD) too. For a given $f(s)$, we can find $\hat{C}_{trt}$ for each trait separately, exactly as shown above. We can then sum the resulting marginal log-likelihoods of each trait to arrive at a composite log-likelihood for $f(s)$ using all traits:

$$LL^{SSD}(f(s)) = LL^{trait\,1}(f(s)) + LL^{trait\,2}(f(s)) + LL^{trait\,3}(f(s)) + \cdots$$

We then maximize $LL^{SSD}(f(s))$ exactly as we would do for a single trait.
Note, that $f(s)$ is parametrized by four parameters. Therefore, for the TSD model we have 5 parameters per trait: $C_{trt}$ plus four parameters for $f(s)$. For the SSD model we have four global parameters for $f(s)$ and one trait-specific parameter, $C_{trt}$. Thus, the SSD is vastly more parsimonious in the number of parameters it uses.

##### 3.8 Adding a penalty to regularize $f(s)$

One complication we discovered is that this inference method misbehaves for very high or low selection coefficients since they do not contribute any GWAS hits. Very low selection coefficients have essentially no causal effect size, i.e. they only produce false positive hits. Very high selection coefficients don't produce common variants at all. Because we condition on variants being hits, the value of $f(s)$ at such extreme selection coefficients doesn't change our likelihood at all. Therefore, without additional information our inference procedure may (and does) assign arbitrarily large weight at these selection coefficients.

To counteract this tendency, we added a small penalty to the likelihood with two terms: one against having target sizes larger than the size of the genome and another against having heritabilities larger than 1. The penalty takes the form

$$-\epsilon\left(\frac{\hat{L}}{3\cdot 10^9}+\frac{\hat{h^2}}{1}\right)$$

per each GWAS hit, where we calculate $\hat{L}$ and $\hat{h^2}$ using $\hat{C}$ and $\hat{f}(s)$ as described above. You can see in fig **??** how with low $\epsilon$ values of $f(s)$ explodes at high and low selection coefficients, while with high $\epsilon$ values of $f(s)$ becomes very narrow. Using our measure of model fit (see below) we find that $\epsilon = 0.01$ regularizes $f(s)$ while maintaining good fit.

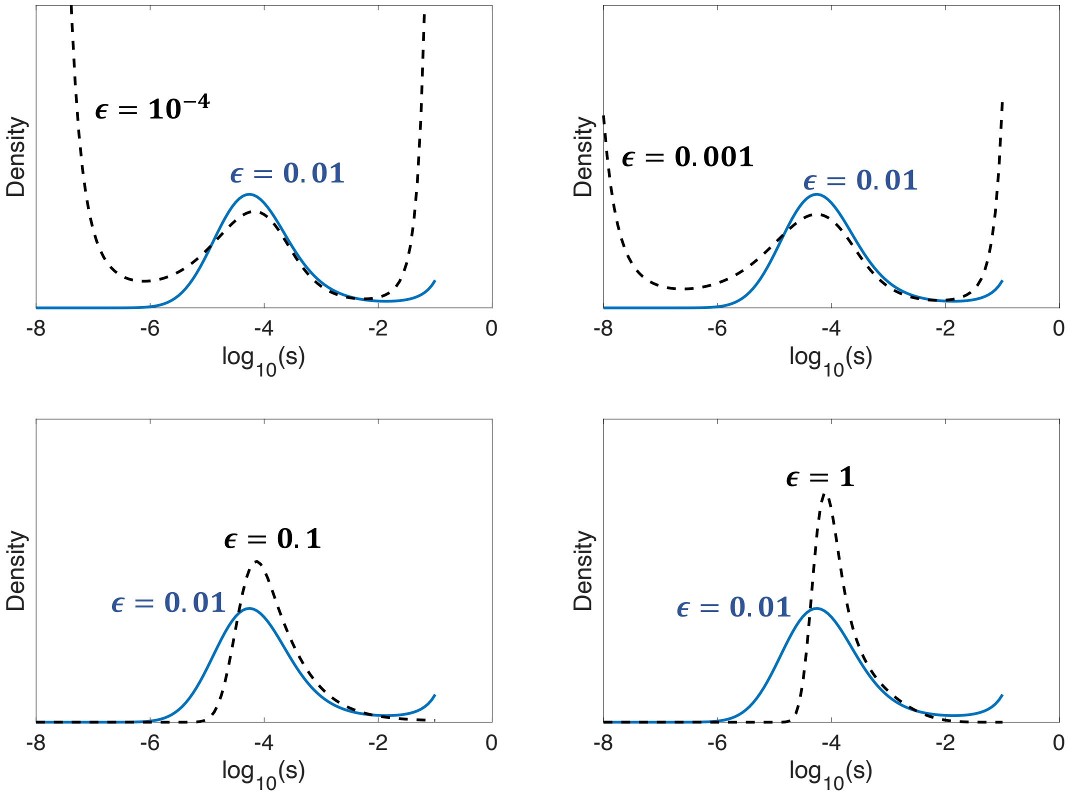

Figure S6: $f(s)$ **for different strengths of regularizing penalty.** *Comparison of the inferred distribution of selection coefficient from our UKbiobank dataset with different values of* $\epsilon$*, the penalty strength. As you can, the distribution in the mid range of selection coefficients is similar for different values of* $\epsilon \leq 0.01$*, but without enough penalty* $f(s)$ *explodes at low and high s values. When the penalty becomes very strong* $f(s)$ *becomes increasingly narrower.*

##### 3.9 Estimating confidence intervals

We estimate confidence intervals by block-wise bootstrap resampling. We use Burisma et al.'s division of the genome into 1702 approximately independent blocks [**?**]. We then resample, with replacement, 1702 blocks, and run the inference on the GWAS hits in the resampled blocks. We repeat this process 100 times to get a distribution the inferred parameters which we then use to estimate confidence intervals.

#### 4 Estimating model fit

Having inferred a model from our data, we are in the need to quantify how well the model fits the data, both each trait and each variant. We introduce a measure that we call the residual p-value – which is a p-value with the inferred model as the null model estimated for each SNP. The residual p-value measures how effects are distributed relative to the model's prediction. We use the distribution of these p-values to estimate model fit to traits, identify outlier variants and set the value of our inference's single hyperparameter.

##### 4.1 The residual p-value

By definition, the one-sided p-value for a measurement $\theta_i$ of a statistic $\theta$ is

$$p_i = Pr(\theta > \theta_i|\text{null})$$

where we know the distribution of $\theta$ under the null. By definition, $p$ in uniformly distributed on the interval (0,1). Therefore, a low p-value allows us to reject the null for measurement $i$.

In GWAS, reported p-values can be thought of as one-sided p-values for the z-score square, $z_i^2$, under a null of no causal effect

$$p_i = Pr(z^2 > z_i^2|\text{null})$$

with $z^2$ having a chi-squared distribution with 1 d.f. under the null.

Our inferred model predicts the distribution of causal effects for variants of a given frequency and so we can define, for a variant of MAF $x_i$ and z-score $z_i$, the residual p-value as

$$p_i = Pr(z^2 > z_i^2|C, f(s), x_i, m_i)$$

with $C$ and $f(s)$ being our estimated model parameters, $x_i$ the MAF and $m_i$ the relative study size. However, our dataset includes only GWAS hits so we need to condition on z-scores being above 5.45. That's easily done under this framework with

$$p_i = \frac{Pr(z^2 > z_i^2|C, f(s), x_i, m_i)}{Pr(z^2 > 5.45^2|C, f(s), x_i, m_i)}.$$

If our model completely describes the co-distribution of MAF and z-score, then for GWAS hits this p-value should be uniformly distributed on the interval (0,1).

##### 4.2 Cross-validated residual p-value

To avoid overfitting bias, we cannot estimate residual p-values on the same data on which we infer model parameters. Instead, we use Burisma et al.'s division of the genome into approximately independent blocks [?]. We then split the genome into 10 parts based on the last digit of the block number. This split ensures that the blocks in each part are well spaced from each other. For each of the 10 parts of the genome, we infer the model on GWAS hits in the other 9 parts. We use the inferred model parameters to calculate the residual p-values for GWAS hits on the held-out part. In this way, we estimate residual p-values for all GWAS hits.

##### 4.3 Measure of model fit for a trait

If our model fits well, we expect the residual p-values for all GWAS hits for a trait to be uniformly distributed. If it does not, we expect to see some deviation. We use the Kolmogorov-Smirnov (KS) p-value for this distribution as our measure of model fit for a trait.

As one can see in Fig. 4 of the main text, these KS p-value are slightly smaller than expected under the null for both the TSD and SSD models. Looking at the KS p-values, no trait stands out as a clear outlier. In fact, with SSD, only 4 out of the 95 traits have KS p-values below the Bonferroni corrected threshold of significance (0.05/95): Mean platelet (thrombocyte) volume, platelet distribution width, glycated haemoglobin, and high light scatter reticulocyte count. Of which, only glycated haemoglobin was also an outlier with TSD. See Supplementary Table 1 in a separate csv file.

##### 4.4 Measure of model fit for variants

We want to identify outlier variants, since they are potentially important to understanding the biology and/or evolution of a trait. We define an outlier p-value as one with

$$p < \frac{0.05}{n_t}$$

with $n_t$ being the number of hits for the trait. This criterion is just an 0.05 threshold with a Bonferroni correction. See list of outliers and our analysis of their biology in Supplementary Table 2.

##### 4.5 Residual p-values for alternative models

We wanted to test the fit of other models to data. We therefore repeated the same procedure, just with other models. We fit a maximum likelihood model where effect sizes are normally distributed with mean zero and variance $A \cdot (x(1-x))^{\alpha}$, with $A$ and $\alpha$ being the model parameters. When we set $\alpha = 0$ and only infer $A$, we get a simple normal distribution and when we infer the value of $\alpha$ this is the "alpha model". We again infer the model on 90% of the genome and calculate residual p-values (conditional on genome-wide significance) for the other 10%.

##### 4.6 Tuning $\epsilon$

We use the residual p-values to tune the hyperparameter controlling the strength of the penalty in our likelihood, $\epsilon$. We tried different values of $\epsilon$, and saw that when $\epsilon < 0.01$ we get unrealistic target sizes and heritabilities. On the other hand, for $\epsilon > 0.01$ the Kolmogorov-Smirnov p-values for model fit for traits become very small, suggesting such large penalty affects model fit. We therefore chose $\epsilon = 0.01$, for this hyperparameter. See Fig. **??**.

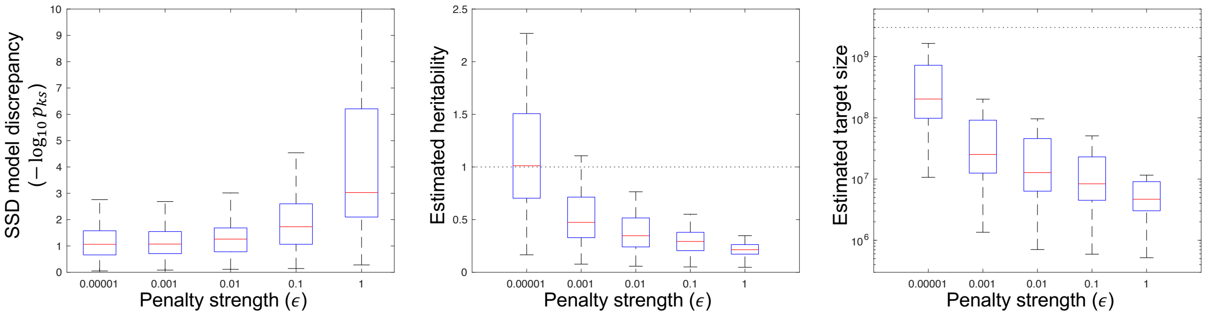

Figure S7: **Tuning the strength of the penalty.** *If the penalty is too strong, the model stops fitting the data. If the penalty is too weak, the estiamted heritabilities and target sizes are unrealistic.*

### 5 Validating our inference using simulations

In order to test our inference, we simulate data from our model and run the inference on it. We use both the SSD model and TSDs. We see that the inference capture the distribution of selection medium coefficients ($-6 < log_{10}(s) < -2$) very well, since they produce GWAS hits. However, contributions to trait architecture from beyond this range are missed, meaning that we may underestimate target sizes and heritabilities. Lastly, we show that if a dataset contains a mixture of traits from different distributions of selection coefficients, then TSDs are a much better fit to the data than the SSD, as expected.

#### 5.1 Simulating a single trait

Given the parameters $L$, $h^2$ and $f(s)$, and in addition study size $M$, we want to simulate a set of hits based on our model with a set of MAFs, z-scores and relative study sizes $\{x_i, z_i, m_i\}$. We use the following algorithm:

1. Set $C = M \cdot \frac{h^2}{L} \frac{1}{E[2q(1-q) \cdot s | f(s)]}$.
2. Draw the number of hits, $\#_{hits}$, from a poisson distribution with mean $L \cdot P(hit|C)$.
3. Calculate the table $P(x, s|C) = \frac{1}{\mathbb{N}} P(hit|C, x, s) P(x|s) f(s)$ for our grid of $x$ and $s$, with $\mathbb{N}$ being a normalization factor such that $\int_s \int_{x>1\%} P(x, s|C) = 1$.
4. Draw $\#_{hits}$ samples from $P(x, s|C)$ to produce the set $\{x_i, s_i\}$.
5. For each sample, draw $m_i$ from $P(m|x)$.
6. For each sample, draw $z_i$ from the distribution $N(0, 1 + C \cdot m_i \cdot s_i \cdot 2x_i(1 - x_i))$ conditional on $|z_i| > 5.45$.

#### 5.2 Simulating sets of traits

We simulate sets of traits, with traits having one of the following 4 specified distributions of selection coefficients (see Fig. **??**):

1. $log_{10}(s) \sim N(-4, 0.5)$
2. $log_{10}(s) \sim N(-3, 0.5)$
3. $log_{10}(s) \sim N(-2, 0.5)$
4. $s \sim \Gamma(0.1, 3 \cdot 10^{-2})$

In each set, we simulate 40 traits with heritabilities drawn uniformly from the interval (0.1,0.9). For a given heritability, we draw the log target size uniformly from the range of target sizes for which the expected number of GWAS hits is above 100. We draw sets from each of the 4 distributions, as well as a set with 10 traits from each distribution. We run both the TSD and SSD models on these sets.

#### 5.3 Validating our inference on simulated traits

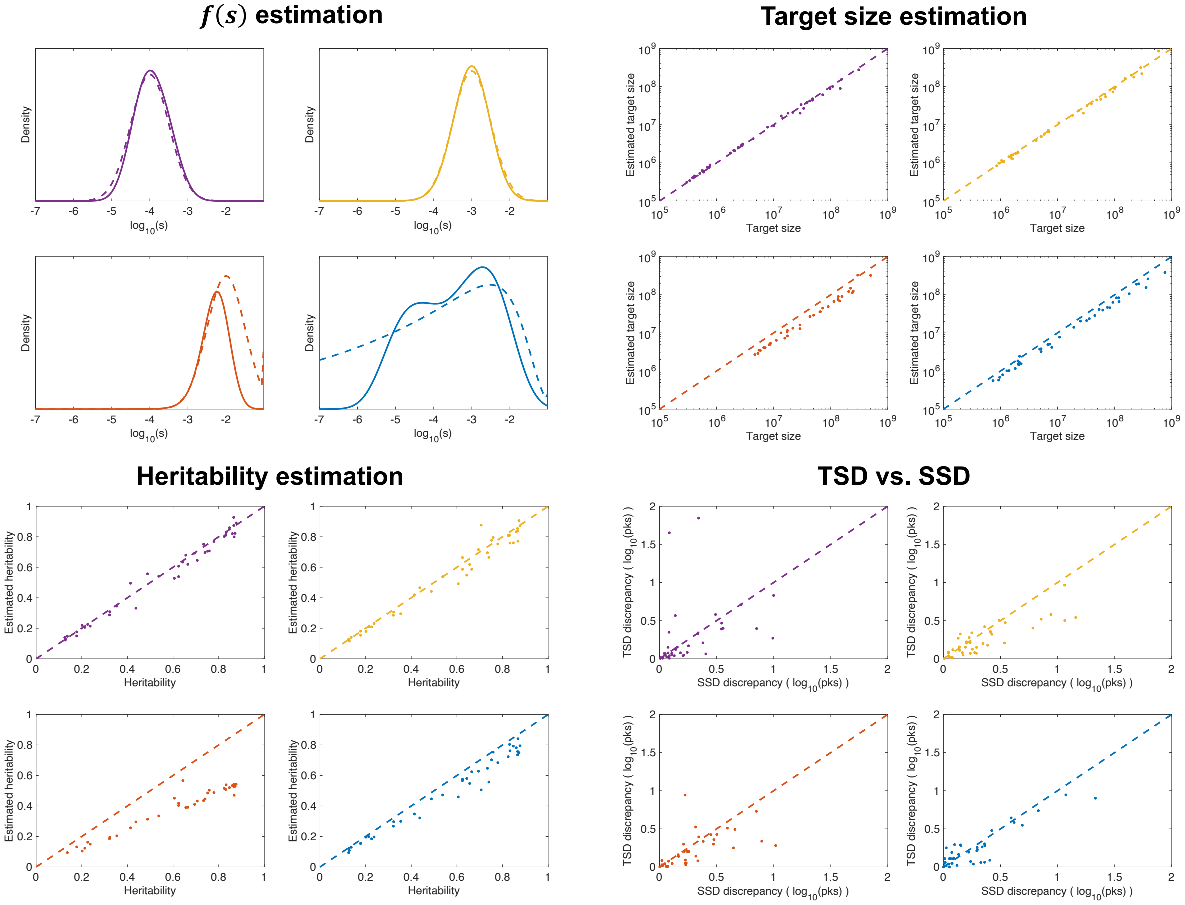

Figure S8: **Inference on simulated datasets.** **(A)** *The distribution of selection coefficients within the range of $-6 < log_{10}(s) < -2$ is well captured by the SSD inference. True distributions in dashed lines, inferred distribution in continuous.* ***(B)*** *Estimates of target size correlate very well with true target size.* ***(C)*** *Estimates of heritability correlate very well with true heritability.* ***(D)*** *TSD and SSD have low and similar discrepancy to data.*

We simulated datasets with study size of $3.6 \cdot 10^5$ and 40 independent traits each. Selection coefficients were drawn from one of 4 distributions of selection coefficients (see Fig. **??**A). For each trait, the heritability was chosen uniformly between 0.1 and 0.9 and the log target size uniformly in the range of target sizes for which the expected number of hits is above 100. We ran our inference (both SSD and TSD) for each such trait.

As you can see in Fig. **??**A, with 40 traits the SSD model captures the distribution of selection coefficients extremely well. However, when the distribution includes significant contributions from high ($log_{10}(s) > -2$) or low ($log_{10}(s) < -6$) selection coefficients, our inference truncates the distribution. Since these extreme selection coefficients do not produce any GWAS hits, our inference puts no weight on them (because we have a parsimony-inducing penalty on our likelihood). Therefore, the distribution of selection coefficients we infer is the distribution at the medium range of selection coefficients ($-6 < log_{10}(s) < -2$).

Our inference only estimates the heritability and target size from within this medium range of selection coefficient. Therefore, for distributions of selection coefficients which are entirely within this range, our inference provides unbiased estimates of the entire heritability and target size, Fig. **??**B&C. However, for distributions with contributions from outside this range, our inference systematically underestimates the heritability and target size by a constant.

As expected, the SSD and TSD have similar levels of discrepancy between inferred model and data, Fig. **??**D. However, when we simulate a dataset with 10 traits from each of the 4 distributions and infer a SSD on it, we see that the inferred distribution only approximates 1 out of the 4 distribution, i.e. it misspecifies the distribution for 30 out of the 40 traits, Figure **??**A. Unsurprisingly, under this scenario, there is huge discrepancy between the SSD model and the data, which is completely absent from the TSD model, Figure **??**B. Compare this result with Fig. 4C of the main text, where we do not see such a difference between TSD and SSD in our data.

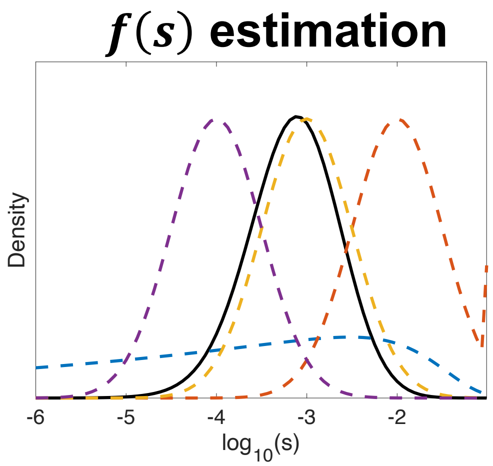

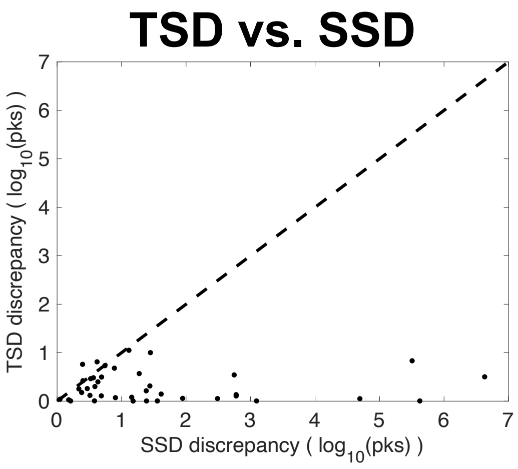

Figure S9: **Inference on a non-SSD dataset.** *Inference on a dataset of 40 traits, 10 from each of 4 different distributions of selection coefficients.* ***(A)*** *The inferred distribution of selection coefficients under a SSD model can match, at most, one of the underlying distributions. True distributions in dashed lines, inferred distribution in a black continuous line.* ***(B)*** *The SSD model has large discrepancy with data, while the TSD model does not.*

### 6 UKBB Dataset

In our inference, we use GWAS hits from 95 continuous traits from the UK biobank. Here we describe the dataset, variant filtering and trait choice.

#### 6.1 GWAS summary statistics for diverse traits

The Neale lab performed standardized GWAS on all available UK biobank phenotypes (`http://www.nealelab.is/uk-biobank`) and we use their summary statistics (version 3). We chose to focus only on continuous phenotypes, using GWAS results based on both sexes and inverse rank-normal transformed ("irnt") phenotypic values. We will address categorical phenotypes and disease risk in future work.

#### 6.2 Trait choice

Based on simulations, we were aiming to keep traits with at least 100 hits. In addition, we wanted to minimize trait repetition and avoid traits whose genetic architecture is concentrated at a single locus.

For trait screening we used PLINK's LD-based clumping [**?**]. For each trait in the GWAS data, we used plink's –clump flag with a p-value threshold of $5 \cdot 10^{-8}$, LD threshold of $r^2 = 0.1$ and physical distance threshold of 1Mb. We only kept traits with at least 100 clumped hits at MAF above 1%. This left us with 138 traits.

Next, if two traits are identical except for handedness, e.g. Arm fat percentage (left) and Arm fat percentage (right), we kept the one with the (slightly) larger number of hits. This left us with 114 traits.

Lastly, we counted the number of hits at each approximately-independent genomic block from [**?**]. If over 10% of hits reside in a single genomic block (out of 1702 blocks) we dropped the trait. At the end of this process, we were left with 96 traits.

#### 6.3 Variant filtering

Starting with the set of 13.7 million variants ascertained by the Neale lab for GWAS ("imputed-v3 Variant QC"), we restricted our analysis to variants passing the following filters: (i) autosomal, (ii) bi-allelic (genomic positions where at most two alleles had frequency ¿0.001), and (iii) MAF¿1%.

Next, we ran COJO [**?**] on each trait to arrive at approximately independent hits and co-estimate their effect sizes. We used the parameters –cojo-p $10^{-6}$ and –cojo-slct. As a reference panel for LD, we used 10000 randomly chosen individuals from the UKBB, passing the following QC measures: (i) reported gender matched the inferred sex from genotype data, (ii) were not heterozygosity outliers,

(iii) did not have excessive number of relatives, and (iv) did not carry sex chromosome aneuploidies. We further restricted the panel to individuals labeled by the UKBB team to be "White British" and were chosen for principal component analysis (PCA) to exclude close relatives.

We filtered out variants with LD score above 300 or INFO score below 0.8. After filtering, the number of hits for one of the traits dropped below 100 and so we were left with the 95 traits in Supplementary Table 1.

#### 6.4 Summaries of GWAS hits

The GWAS hits for the 95 traits differ in their number, effect sizes and minor allele frequencies, as seen in Figure ??. Despite restricting ourselves to traits with at least 100 hits, the number of hits still spans an order of magnitude (Fig. ??A). These hits vary in their z-scores, and the magnitude of z-scores doesn't have any clear relation to the number of hits (Fig. ??A). However, the mean MAF is very similar across all trait (Fig. ??B), which might be the result of restricting ourselves only to common variants. Since the number of hits and their effect on traits vary between traits, the proportion of phenotypic variance explained by the hits for each trait varies greatly (Fig. ??C).

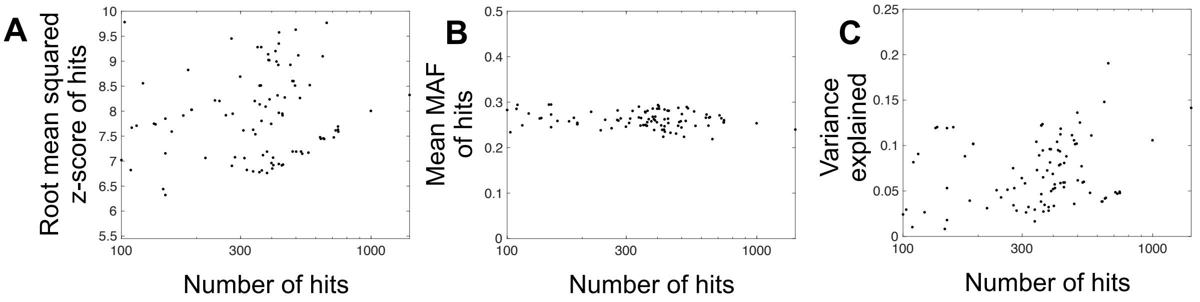

Figure S10: **Summaries of GWAS hits for the 95 traits.** *(A) The root mean squared z-differs between traits and is not determined by the number of hits, which spans an order of magnitude. (B) In contrast, the mean MAF is very similar across all traits. (C) The proportion of phenotypic variance explained by hits also greatly differs between traits.*

#### 6.5 Functional followup of outlier variants

16 variants in our dataset are clear outliers, having significantly low residual p-values (see Section ??). In order to understand which genes they affect we annotated each SNP's gene and variant type using dbSNP. The majority of variants were within gene boundaries of a known gene and were annotated with that gene. Two variants (rs700750 and rs386698213) were not within a gene window and thus were annotated as noncoding variants. We annotated these noncoding variant's with their nearest genes using a list of all genes in any GO, KEGG, or Reactome MSigDB pathway (in order to avoid pseudogenes and genes of unknown function) and manual curation using GeneCards to prioritize a single gene per SNP. GeneCards was also used to identify each gene's known function for all 16 variants. GWAS catalog was used to identify each SNP's relevant trait and disease associations. These results are presented in Supplementary Table 2 where candidate mechanisms for how each variant affects their respective gene, protein, and the trait are listed in column 11.

##### 6.6 Computational reduction in study size

When compare the genetic architecture of traits, we want to equate their median study sizes. In order to so, we take the output of COJO, which has a p-value threshold of $10^{-6}$. We introduce additional Gaussian noise to each z-score:

$$z_{330k} = \sqrt{\frac{330,000}{M_{med}}} \cdot z + \sqrt{1 - \frac{330,000}{M_{med}}} \cdot \epsilon$$

with $M_{med}$ being the median study size for the trait and $\epsilon$ drawn from a Normal distribution with mean 0 and variance 1.

#### 7 Allele ages

In this section, we use the output of RELATE estimation of genome-wide genealogies [**?**] to show that GWAS hits are younger than matched alleles. While the distribution of allele ages of matched alleles matches our prediction for neutral alleles, the distribution of allele ages for GWAS hits deviates from our predictions. We explain why this deviation is due to a bias in the point estimator for allele ages for alleles under selection. We use simple timescale arguments to suggest a heuristic correction for this bias that resolves this discrepancy.

##### 7.1 RELATE output

At each genomic location, RELATE has estimated the local genealogy among the 1,000 genome samples, spanning a diverse set of global populations. These genealogies are mainly informed by local haplotype structure.

Each variant is then placed on this local genealogy and RELATE reports the branch in the genealogy on which this variant has arisen. For this terminal branch, RELATE reports estimates of the branch's beginning and end, which provide a lower an upper bound on that variant's age. We denote these bounds as $T_{low}$ and $T_{up}$, see Fig. **??**A.

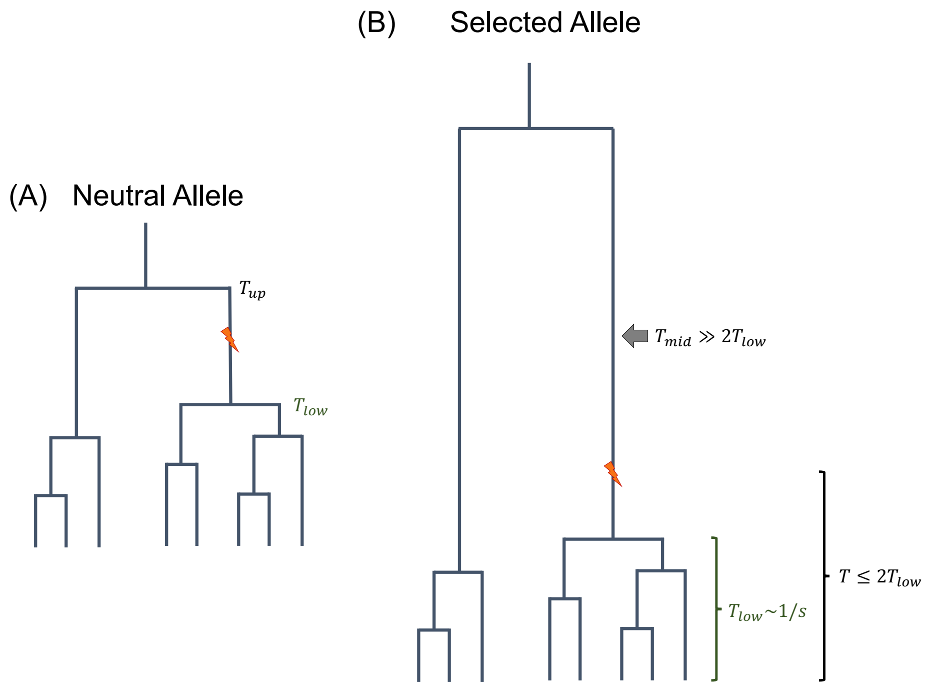

Figure S11: **Schematic of mutation origin on terminal branch.** ***(A)*** *By placing a mutation on the genealogy inferred at a locus we can bound the mutation age from above and below by the edges of the branch on which it arose.* ***(B)*** *For an allele under selection, a mutation could only have arisen in the lower part of the branch.*

#### 7.2 GWAS hits are younger than matched controls

RELATE documentation recommends treating the midpoint of the branch on which a variant rose as its age, i.e. $\hat{T}_{mid} = \frac{1}{2}(T_{low} + T_{up})$. We used this estimator to estimate the age of GWAS hits for all 95 traits in our dataset. To see if they are indeed younger than neutral alleles, we matched each hit with a variant of a similar derived allele frequency. We chose the matched SNPs from regions of low background selection (top 10% of B statistic, taken from Murphy et al [?]). We see that indeed GWAS hits are younger than matched alleles (Fig. **??**).

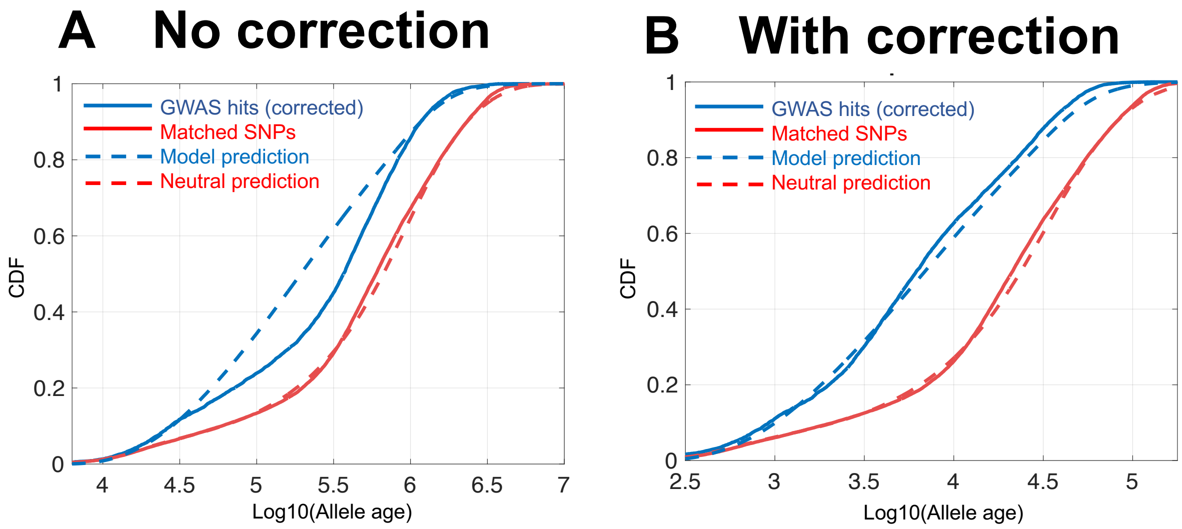

Figure S12: **Distribution of allele ages.** ***(A)*** *GWAS hits are younger than matched alleles. While the distribution of allele ages of matched allele ages aligns well with neutral prediction, there is some discrepancy between our prediction for GWAS hits and RELATE based estimates.* ***(B)*** *After applying our heuristic correction, this discrepancy disappears.*

#### 7.3 Allele age prediction

We next wanted to see if the distribution of allele ages matches our predictions so we ran simulations, under our demographic model, for different selection coefficients. For each variant, we recorded its frequency and age. Then, for each selection coefficient and frequency bin we could estimate the distribution of allele ages $P(T|x, s)$.

As a first step, we show that our predictions for neutral alleles match what we see for matched SNPs. For neutral predictions, we estimate the distribution of allele ages for variants with the same MAFs as our GWAS hits. We use $s = 10^{-6}$ as a proxy for neutrality. For GWAS hit $i$, we have a distribution of age conditional on its frequency $P(T|x_i, s = 10^{-6})$. The overall distribution is just an average over the distributions for each SNP

$$p_{neut}(T) = \frac{1}{n} \sum_i P(T|x_i, s = 10^{-6})$$

with $n$ being the total number of hits. Our prediction for neutral alleles closely matches the distribution of allele ages produced by the midbranch point estimator $T_{mid}$, see Fig. **??**.

Next, we want to compare the predictions of our inferred model to the observed distribution. For our model predictions of allele age for GWAS hits, it's easier to think first of a single trait. We have a distribution of selection coefficients $f(s)$ and a distribution of allele frequencies $P(x|s)$. However, the probability of an allele being a hit depends on its allele frequency, as discussed above, and we denote it as $P(hit|C, x, s)$. $C$ is the power parameter which we estimate for each trait and is proportional to $h^2/L$. The distribution of allele ages for hits for this trait is then

$$P(T|C) = \frac{\int_x \int_s P(T|x, s) P(hit|C, x, s) P(x|s) f(s)}{P\left(hit|C, f(s)\right)}.$$

The distribution for hits of all traits is just a weighted sum over the single trait distribution -

$$P(T) = \frac{\sum_t n_t \cdot P(T|C_t)}{\sum_t n_t}$$

with $C_t$ being the scaling factor for trait $t$, $n_t$ the number of hits for the trait, and the sum running over all traits. We calculated this distribution numerically. It wasn't fun.

This prediction does not fully align with what we observe, see Fig. **??**.

#### 7.4 Bias in allele age estimation

RELATE gives us an estimate of the two edges of the terminal branch on which an allele arose in the genetic genealogy. For a neutral allele, RELATE documentation suggests the midpoint of this branch as an estimator of allele age. However, this makes little sense for common alleles under selection.

Many GWAS hits originate in long branches, with the branch midpoint more than twice the branch starting point $T_{low}$, see Fig. ??B. Such branches exist due to a combination of the out-of-Africa bottleneck, population structure within Africa before the bottleneck and the limited resolution the 1000 genomes dataset gives for the genetic genealogy before the out-of-Africa exodus.

Unlike a neutral allele, a selected allele could not have arisen at any point on the branch. After arising by mutation, a selected variant either goes extinct after a few generations or rises in frequency on a timescale of $1/s$ generations. Regardless of the mode and direction of selection, selected variants have sojourn times of the order of $1/s$. The age of such a variant is then smaller (in order of magnitude) then $T_{low} + 1/s$. Assuming the variant is presently common and present in multiple samples in a dataset, the time until all copies coalesce, $T_{low}$, is also of the order of $1/s$. The conclusion is that the age of such a variant is smaller then $2 \cdot T_{low}$ (at least in order of magnitude ). That is, when a selected alleles maps to a long branch, it must have arisen at the lower part of the branch, see Fig. ??B.

#### 7.5 Heuristic point estimate of allele age

We found that a simple heuristic correction solves the bias. We sought a point estimate of allele ages, $\hat{T}$, that satisfies $\hat{T} - T_{low} \leq T_{low}$. The simplest estimator is just to take $\hat{T} = \min(\hat{T}_{mid}, 2 \cdot T_{low})$. This estimator resolves the discrepancy, see Fig. ?? and Fig. 5 in the main text.

We did however test a whole family of estimators of the form $\hat{T}_\lambda = \min(\hat{T}_{mid}, \lambda \cdot T_{low})$, such that $\hat{T}_2$ is the estimator we use in the main text and we have the limits $\hat{T}_1 = T_{low}$ and $\hat{T}_\infty = T_{mid}$. In Fig. ??, you can see that our estimates are not sensitive to the exact choice of $\lambda$, with all $1.5 < \lambda < 4$ giving similar results. If $\lambda$ is too small, the estimator becomes insensitive to $T_{high}$ and allele ages are downwardly biased. If $\lambda$ is too big, long branches create a bias in allele ages.

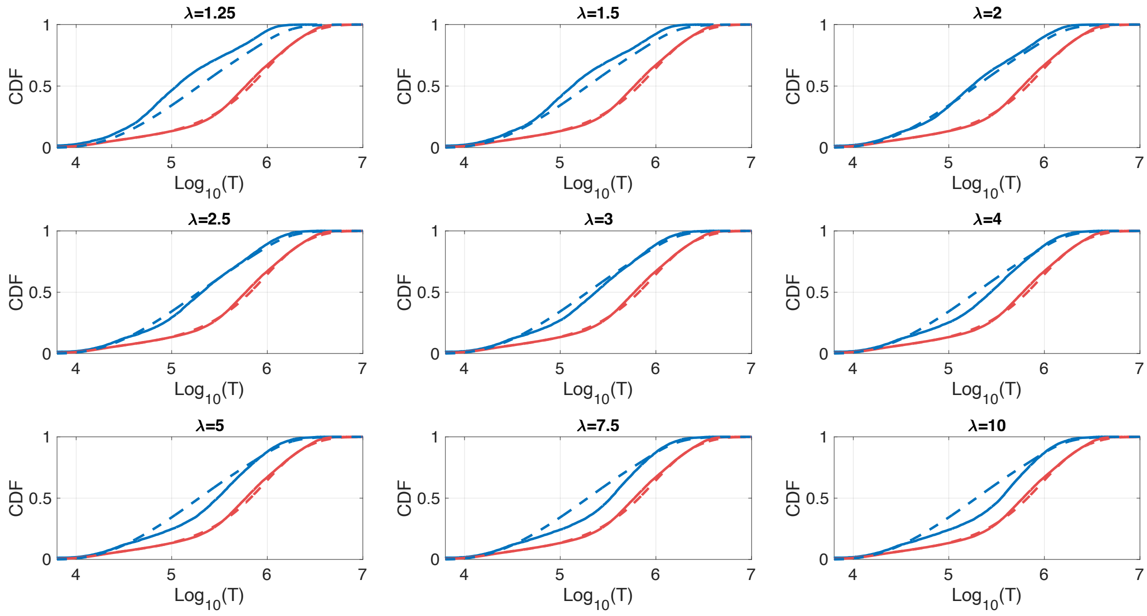

Figure S13: **Heuristic estimators of allele ages.** *We tried different variations of our heuristic estimator and we see that for a wide range of $\lambda$ values the distributions of allele ages closely matches our prediction.*

##### 7.6 Sensitivity to estimates of $s$

To test how sensitive our predictions are to the inferred distribution of selection coefficients, we tested the accuracy of our predictions when $f(s)$ is shifted up or down by $10^{0.5}$ (Fig. **??**A). As you can see in Fig. **??**B, these shifts in selection coefficient result in shifts in the distribution of allele ages such that the predicted distributions no longer match the observed distribution.

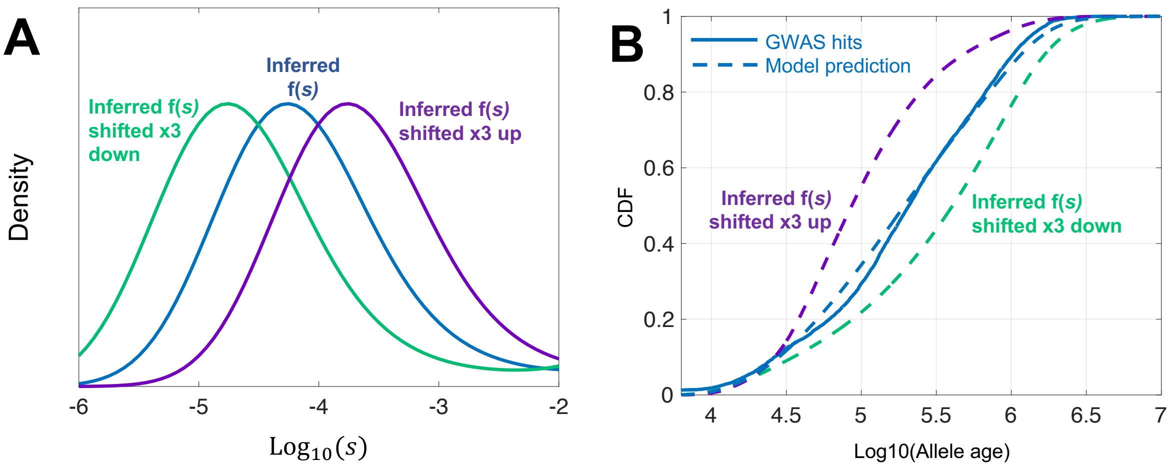

Figure S14: **Prediction of allele ages with shifted distributions of selection coefficients.** *(A) We take the inferred distribution of selection coefficients under the SDS model and shift it up (in purple) or down (in green) by* $10^{0.5}$. *(B) The resulting predicted allele ages are shifted and no longer match the observed distribution of allele ages for GWAS hits.*

#### 8 Similarities in genetic architectures after scaling

In this section, we provide figures analogous to Fig. 7 of the main text, but only for effect sizes. We look at all traits with median study size above $3.3 \cdot 10^5$. We focus on one trait at each row and plot in blue the CDF of effect sizes for that trait. In grey, we plot the CDF of effect sizes for all other traits with $h^2/L$ bigger than that of the focal trait.

We then scale the effect sizes and plot the CDF of the scaled effect sizes for the focal trait (in blue) and all the other traits with $h^2/L$ bigger than that of the focal trait together (in grey). We join the signal for all traits together to reduce noise, since after scaling and thresholding we are left with only a few hits for some traits.

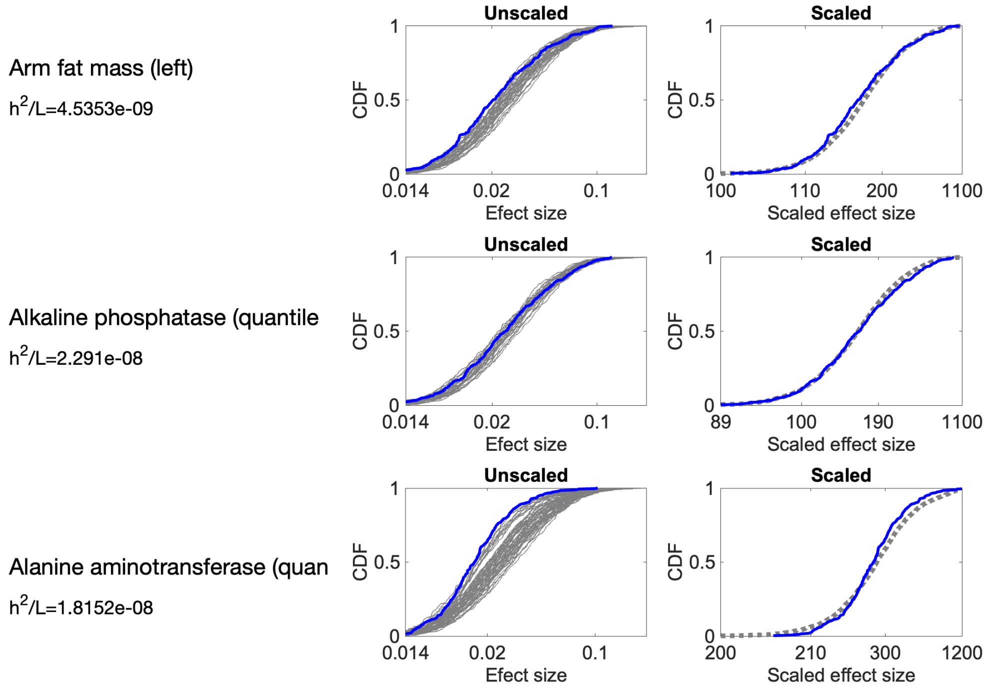

Arm fat mass (left)
$h^2/L$=4.5353e-09
Alkaline phosphatase (quantile
$h^2/L$=2.291e-08
Alanine aminotransferase (quan
$h^2/L$=1.8152e-08
Unscaled
Scaled
CDF
0
0.5
1
0.014
0.02
0.1
Efect size
100
110
200
1100
89
190
200
210
300
1200
Scaled effect size

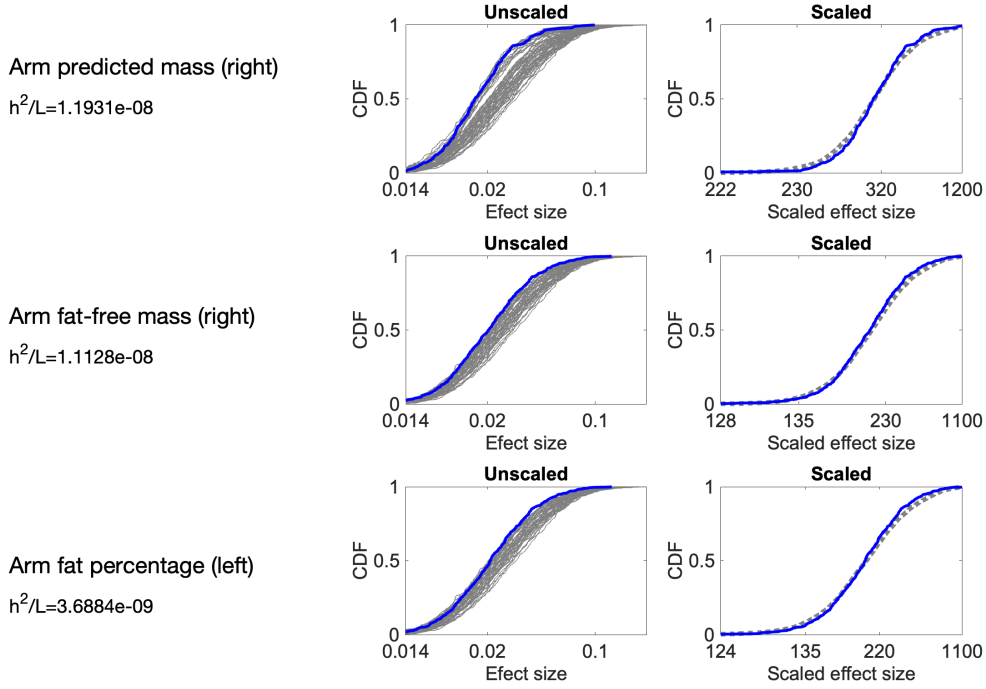

Arm predicted mass (right)
$h^2/L$=1.1931e-08
Arm fat-free mass (right)
$h^2/L$=1.1128e-08
Arm fat percentage (left)
$h^2/L$=3.6884e-09
Unscaled
Scaled
CDF
0
0.5
1
0.014
0.02
0.1
Efect size
222
230
320
1200
128
135
1100
124
220
Scaled effect size

Body fat percentage

$h^2/L$=3.8581e-09

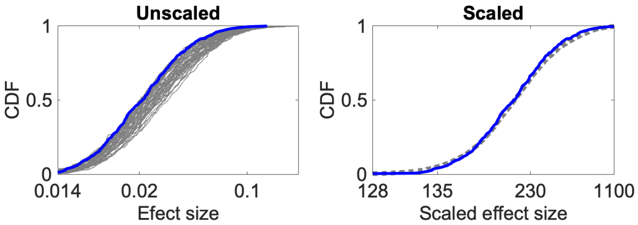

Basophill percentage

$h^2/L$=3.1742e-08

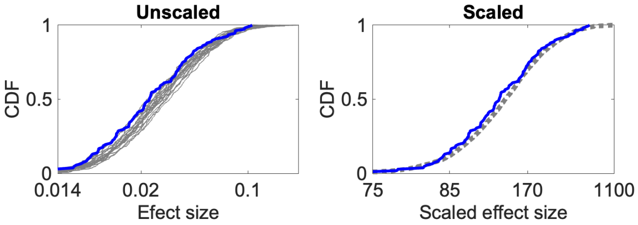

Basal metabolic rate

$h^2/L$=1.1125e-08

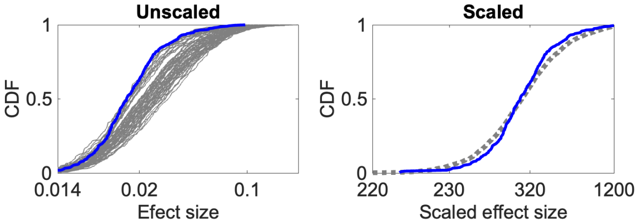

Creatinine (quantile)

$h^2/L$=1.8166e-08

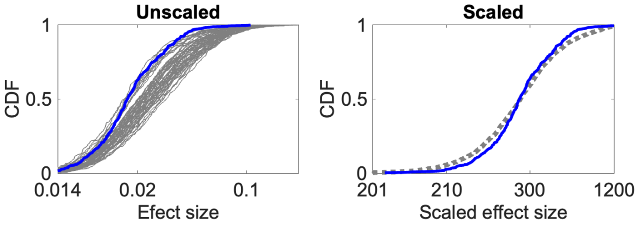

Body mass index (BMI)

$h^2/L$=4.3304e-09

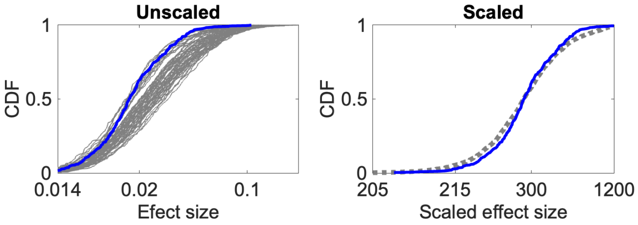

Body mass index (BMI)

$h^2/L$=4.4672e-09

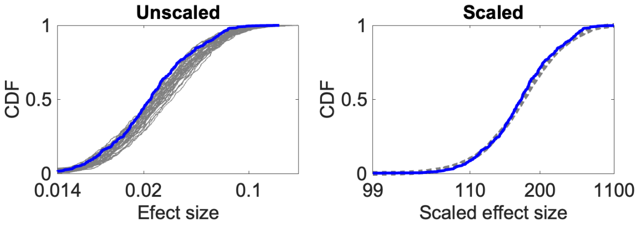

Gamma glutamyltransferase (qua

$h^2/L$=4.3597e-08

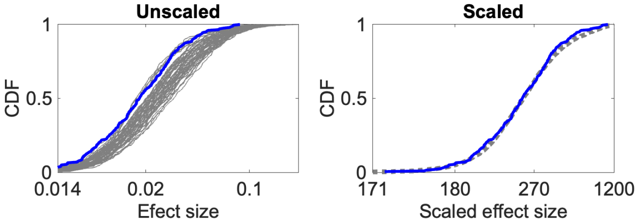

Eosinophill percentage

$h^2/L$=3.7344e-08

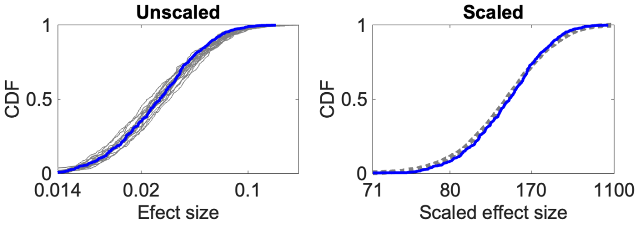

Diastolic blood pressure, auto

$h^2/L$=6.5176e-09

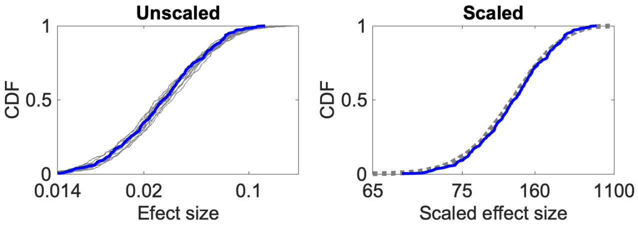

Haemoglobin concentration

$h^2/L$=1.9585e-08

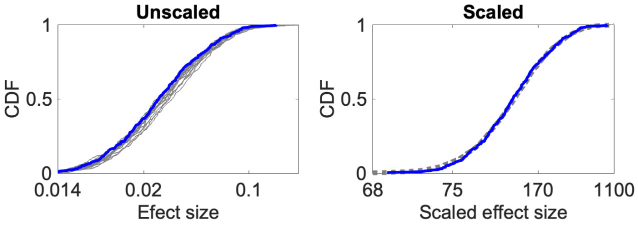

Haematocrit percentage

$h^2/L$=1.8258e-08

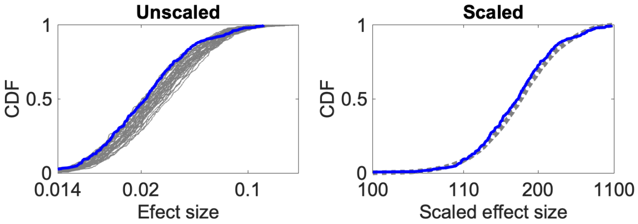

Glycated haemoglobin (quantile

$h^2/L$=3.8834e-08

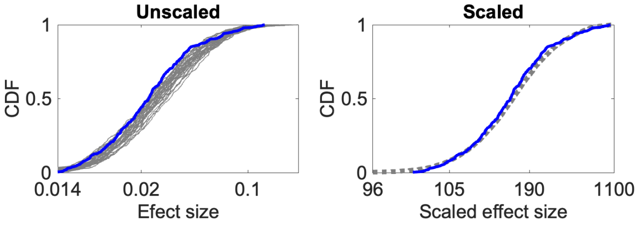

High light scatter reticulocyt

$h^2/L$=3.9676e-08

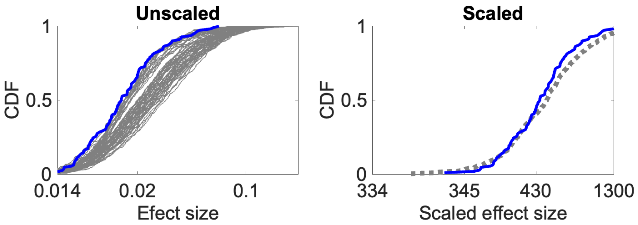

High light scatter reticulocyt

$h^2/L$=3.8779e-08

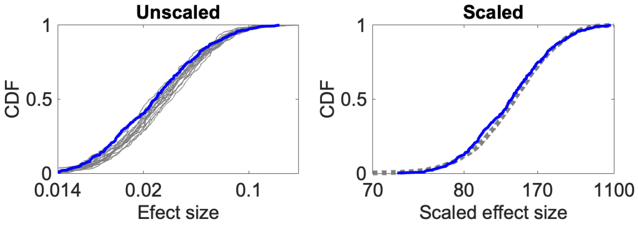

Hand grip strength (right)

$h^2/L$=1.6597e-09

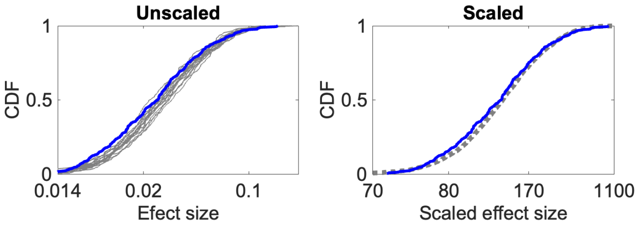

Immature reticulocyte fraction

$h^2/L$=3.9688e-08

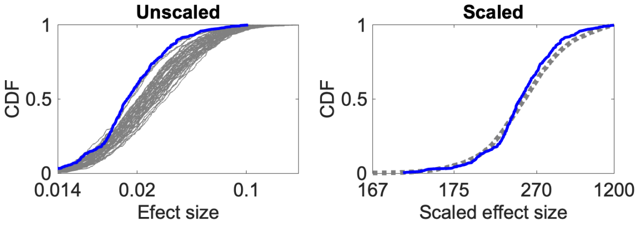

IGF-1 (quantile)

$h^2/L$=2.3527e-08

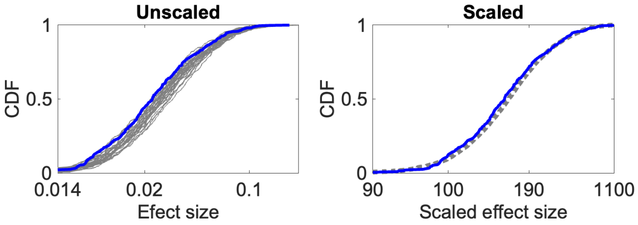

Hip circumference

$h^2/L$=6.7477e-09

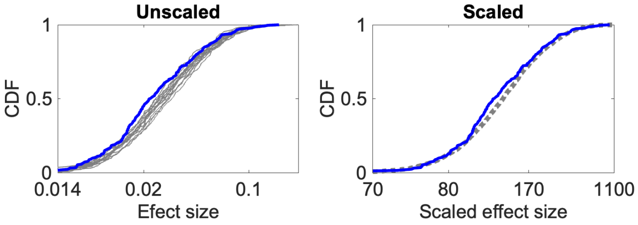

High light scatter reticulocyt

$h^2/L$=3.9676e-08

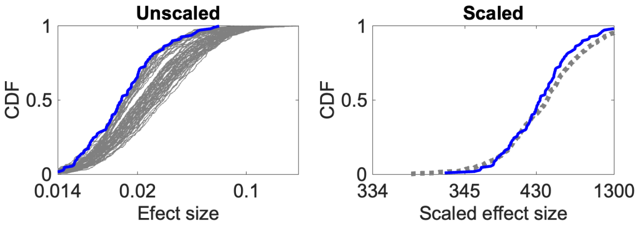

High light scatter reticulocyt

$h^2/L$=3.8779e-08

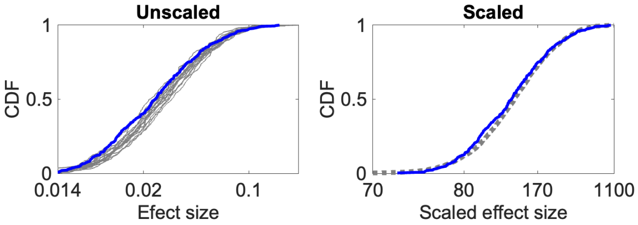

Hand grip strength (right)

$h^2/L$=1.6597e-09

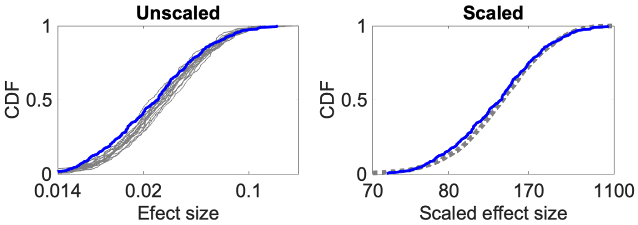

Impedance of whole body

$h^2/L$=7.35e-09

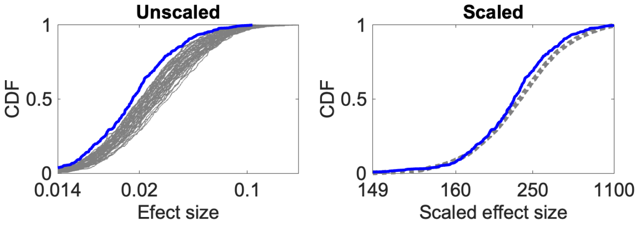

Impedance of leg (left)

$h^2/L$=6.5544e-09

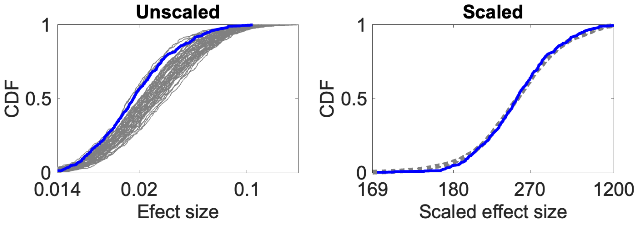

Impedance of arm (left)

$h^2/L$=8.0629e-09

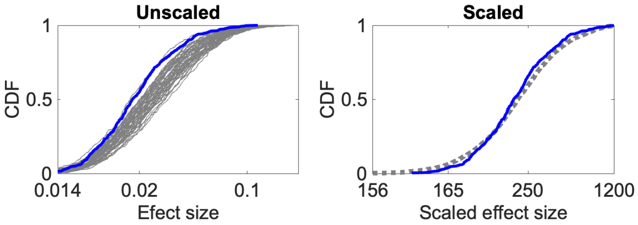

Leg fat-free mass (right)

$h^2/L$=1.1644e-08

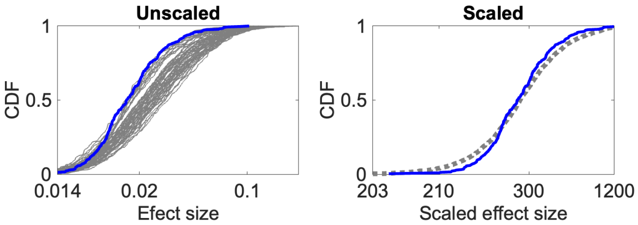

Leg fat percentage (left)

$h^2/L$=3.9472e-09

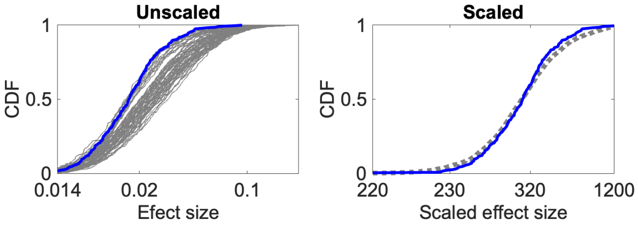

Leg fat mass (left)

$h^2/L$=4.5337e-09

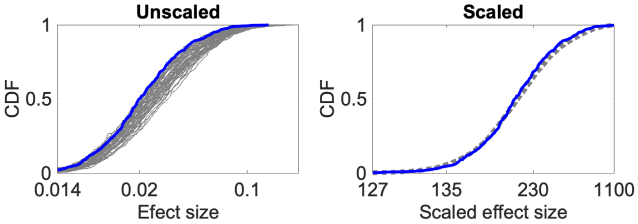

Lymphocyte percentage

$h^2/L$=1.916e-08

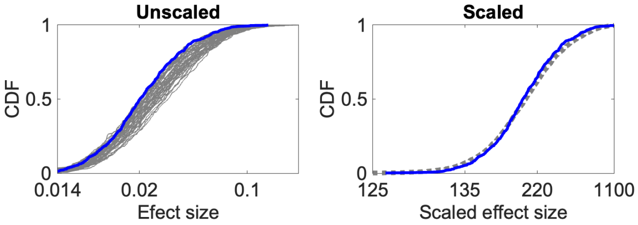

Lymphocyte count

$h^2/L$=2.2002e-08

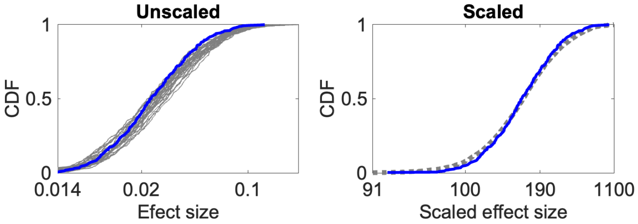

Leg predicted mass (right)

$h^2/L$=1.1648e-08

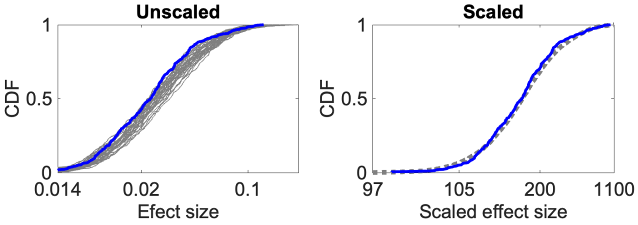

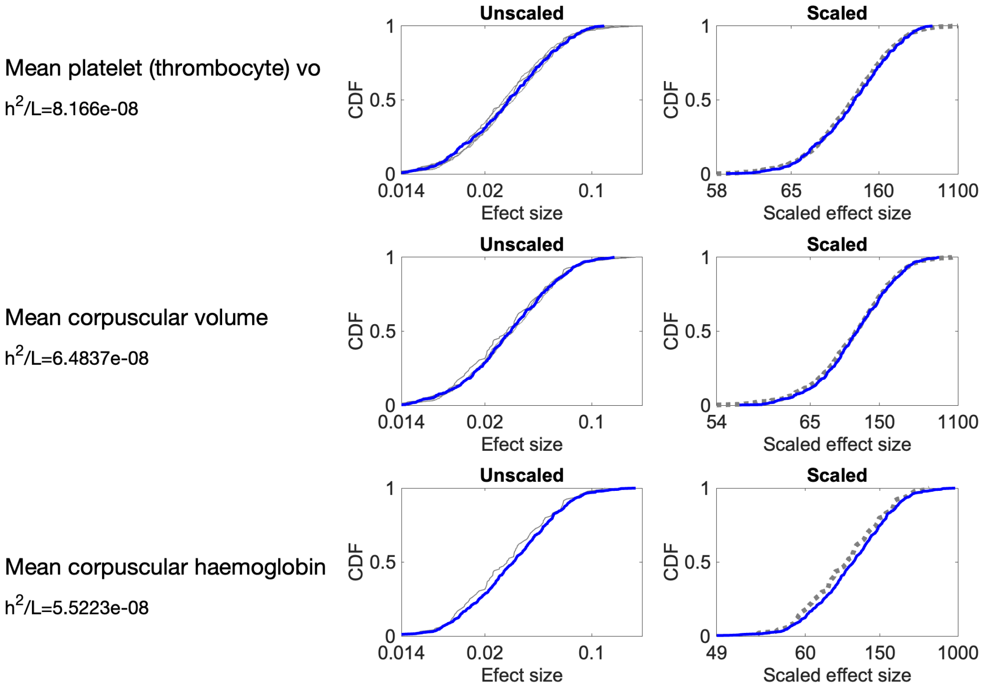
Mean platelet (thrombocyte) vo
h2/L=8.166e-08
Mean corpuscular volume
h2/L=6.4837e-08
Mean corpuscular haemoglobin
h2/L=5.5223e-08
Unscaled
Scaled
CDF
Efect size
Scaled effect size
0.014
0.02
0.1
58
65
160
1100
54
150
49
60
1000

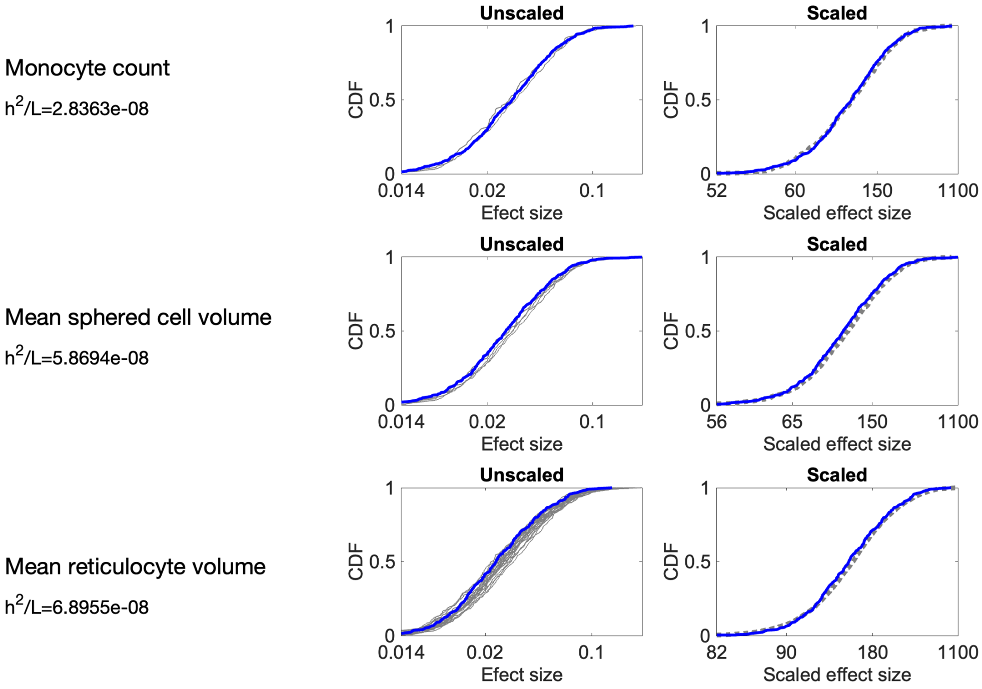
Monocyte count
h2/L=2.8363e-08
Mean sphered cell volume
h2/L=5.8694e-08
Mean reticulocyte volume
h2/L=6.8955e-08
Unscaled
Scaled
CDF
Efect size
Scaled effect size
0.014
0.02
0.1
52
60
150
1100
56
65
82
90
180

Neutrophill percentage

$h^2/L$=1.5221e-08

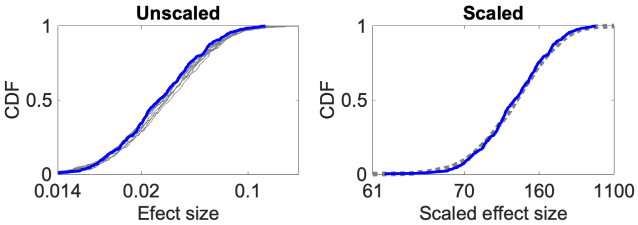

Neutrophill count

$h^2/L$=2.1001e-08

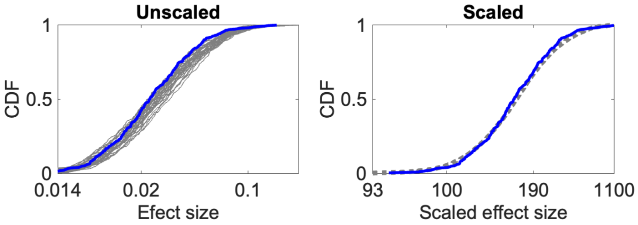

Monocyte percentage

$h^2/L$=4.9267e-08

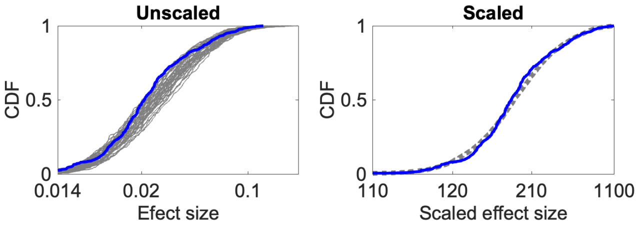

Platelet distribution width

$h^2/L$=5.2644e-08

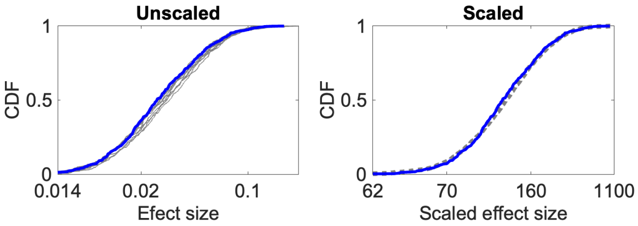

Platelet crit

$h^2/L$=3.1724e-08

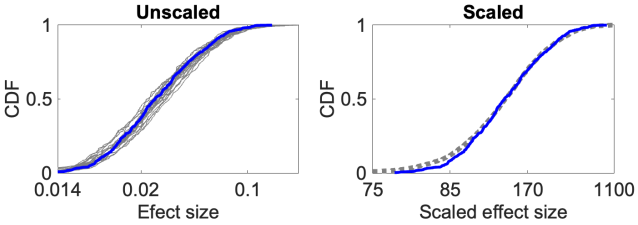

Platelet count

$h^2/L$=4.8039e-08

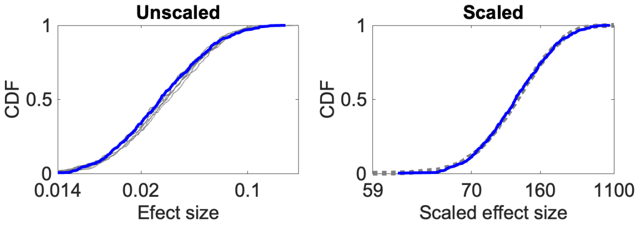

Red blood cell (erythrocyte) d

$h^2/L$=5.3868e-08

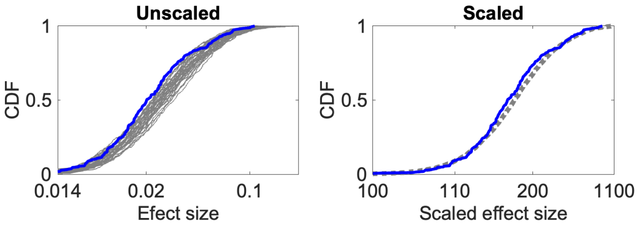

Red blood cell (erythrocyte) c

$h^2/L$=2.7041e-08

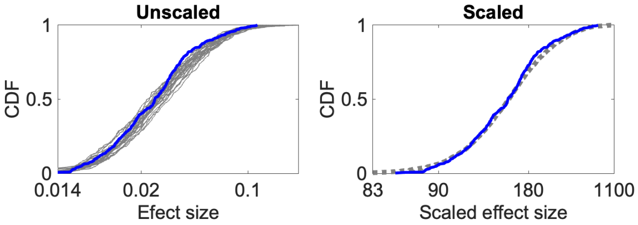

Pulse rate, automated reading

$h^2/L$=1.7989e-08

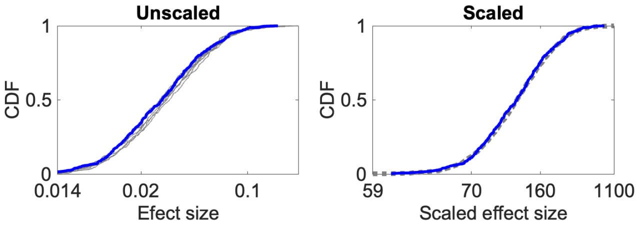

Sitting height

$h^2/L$=2.1888e-08

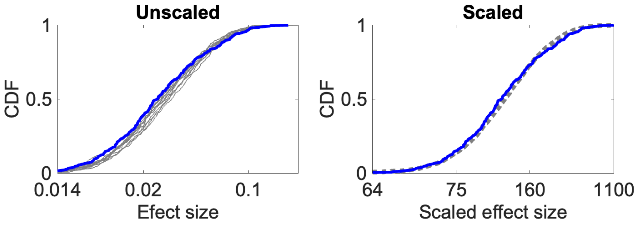

Reticulocyte percentage

$h^2/L$=4.7718e-08

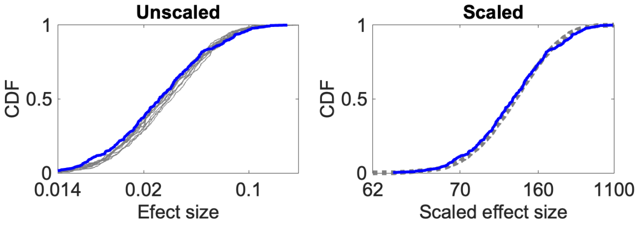

Reticulocyte count

$h^2/L$=4.6645e-08

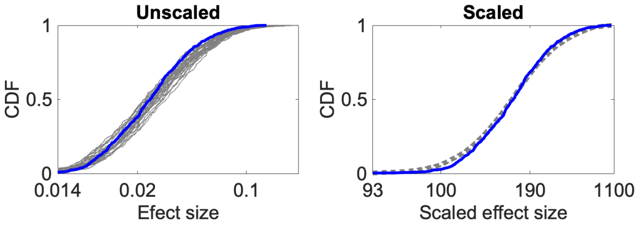

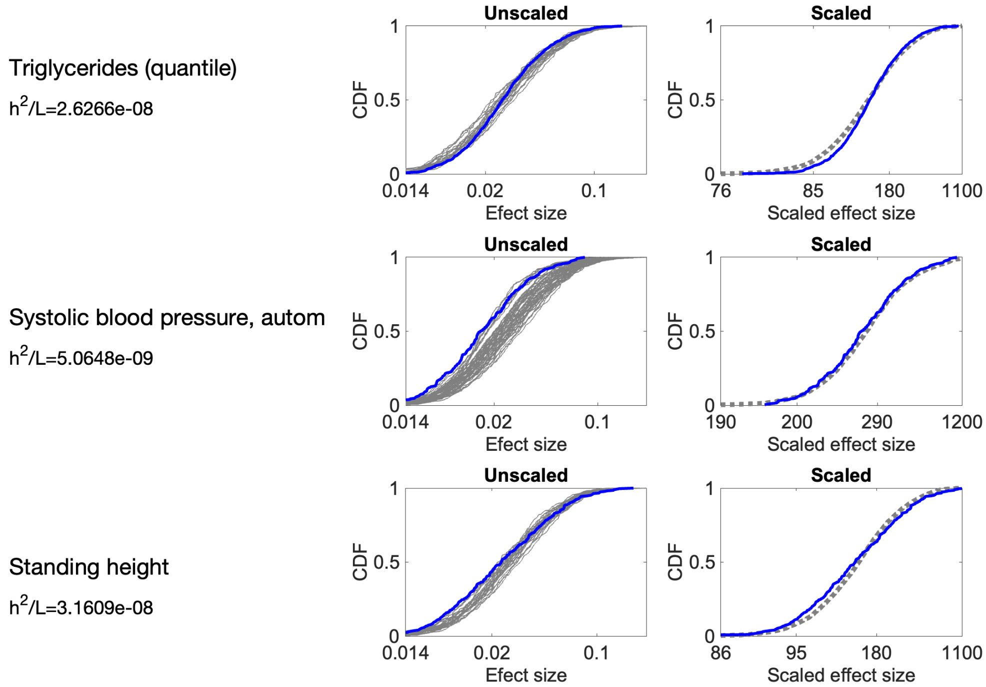

Triglycerides (quantile)
$h^2/L$=2.6266e-08
Systolic blood pressure, autom
$h^2/L$=5.0648e-09
Standing height
$h^2/L$=3.1609e-08
Unscaled
Scaled
CDF
Efect size
Scaled effect size
0.014
0.02
0.1
76
85
180
1100
190
200
290
1200
86
95

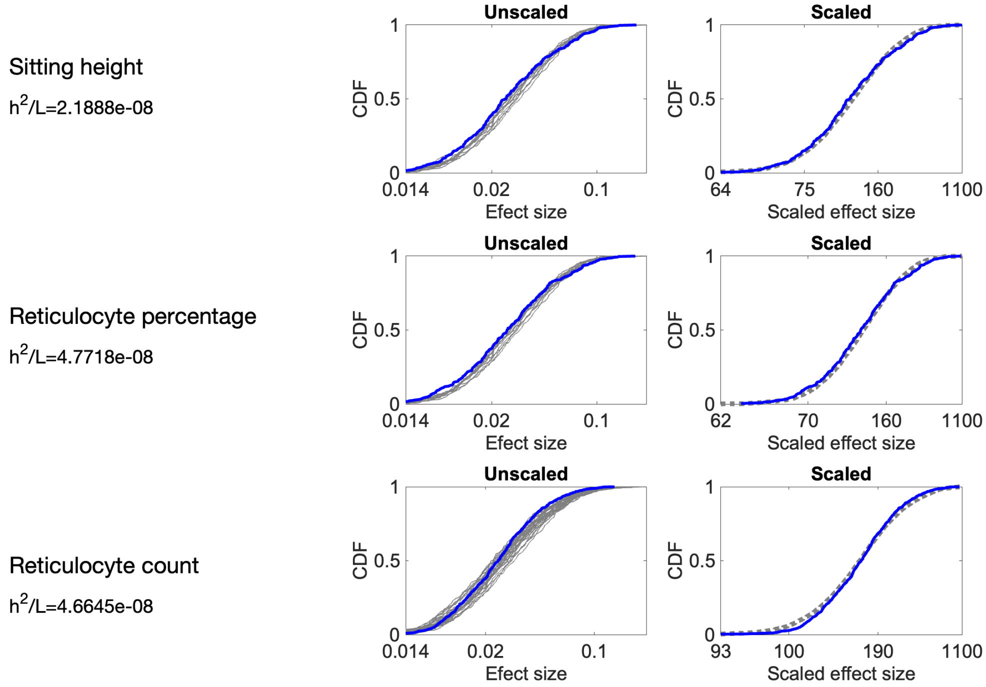

Sitting height
$h^2/L$=2.1888e-08
Reticulocyte percentage
$h^2/L$=4.7718e-08
Reticulocyte count
$h^2/L$=4.6645e-08
Unscaled
Scaled
CDF
Efect size
Scaled effect size
0.014
0.02
0.1
64
75
160
1100
62
70
93
100
190

Trunk fat-free mass

$h^2/L$=1.5731e-08

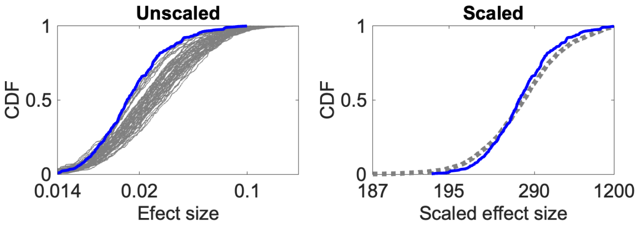

Trunk fat percentage

$h^2/L$=3.8603e-09

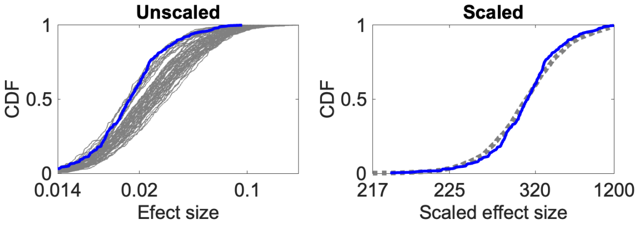

Trunk fat mass

$h^2/L$=5.207e-09

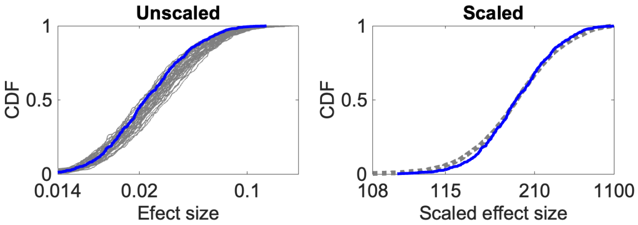

Waist circumference

$h^2/L$=3.543e-09

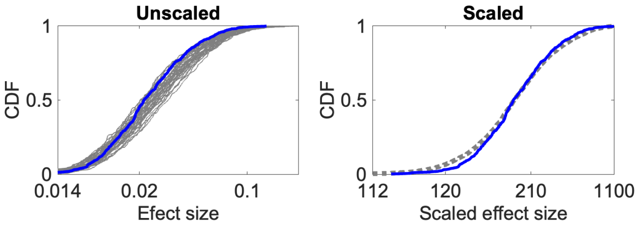

Urea (quantile)

$h^2/L$=3.9763e-08

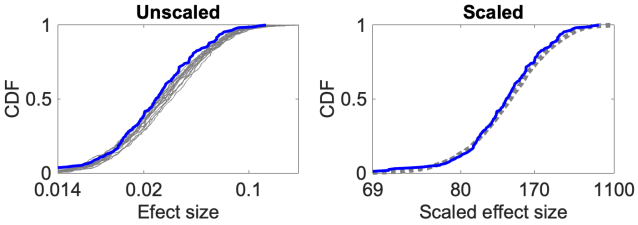

Trunk predicted mass

$h^2/L$=1.4342e-08

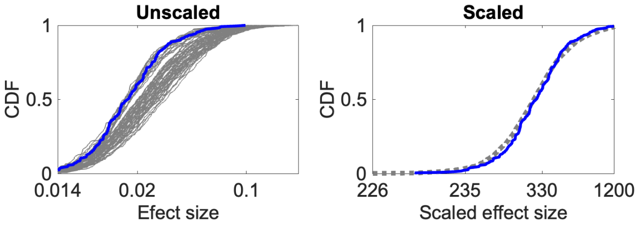

White blood cell (leukocyte) c

$h^2/L$=1.8256e-08

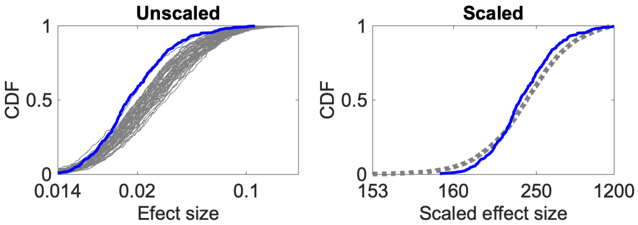

Weight

$h^2/L$=7.3475e-09

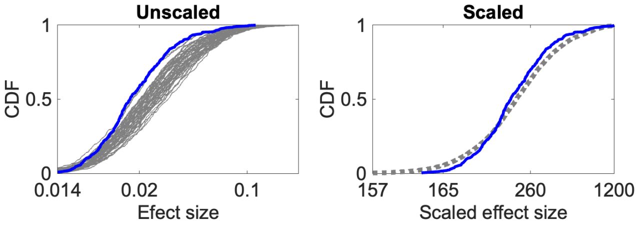

Weight

$h^2/L$=7.7589e-09

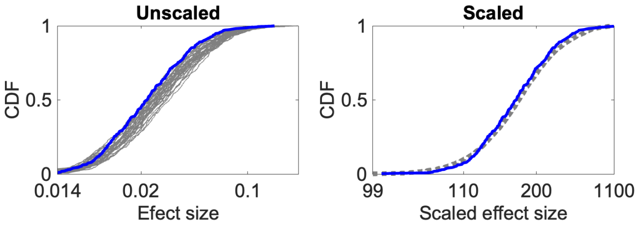

Whole body water mass

$h^2/L$=1.4e-08

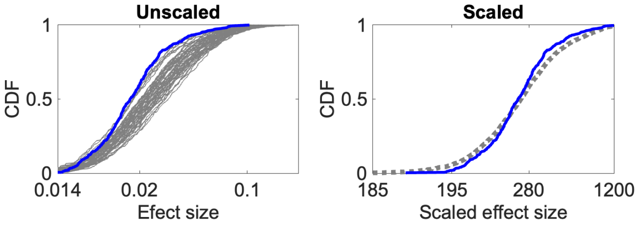

Whole body fat-free mass

$h^2/L$=1.3376e-08

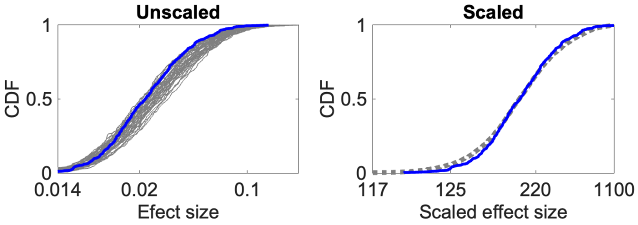

Whole body fat mass

$h^2/L$=5.3344e-09

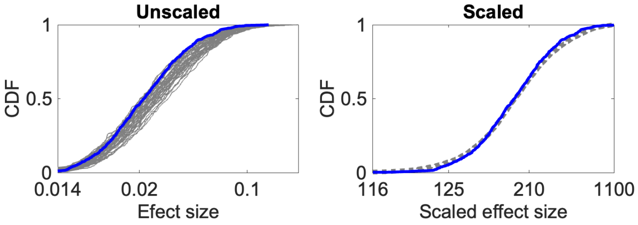

### 9 Supplementary tables

#### 9.1 Table S1: Inferred target size and heritability for all 95 traits

Table is attached as a separate csv file.

#### 9.2 Table S2: Outlier SNPs

| CHR | BP | Traits | Gene | rsid | Type of Mutation |
|---|---|---|---|---|---|
| 1 | 248039451 | Immature reticulocyte fraction | TRIM58 | rs3811444 | missense |
| 2 | 27730940 | Triglycerides (quantile) | GCKR | rs1260326 | missense |
| 6 | 26104632 | Haemoglobin concentration | SLC17A2 | rs386698213 | noncoding |
| 7 | 46753491 | IGF-1 (quantile) | IGFBP3 | rs700750 | noncoding |
| 10 | 71094504 | Haematocrit percentage<br>Haemoglobin concentration | HK1 | rs17476364 | intron |
| 14 | 94844947 | Albumin (quantile) | SERPINA1 | rs28929474 | missense |
| 16 | 53802494 | Arm fat mass (left)<br>Arm fat percentage (left)<br>Body fat percentage<br>Body mass index (BMI)<br>Hip circumference<br>Impedance of leg (left)<br>Impedance of whole body<br>Leg fat mass (left)<br>Leg fat percentage (left)<br>Trunk fat mass<br>Waist cirumference<br>Weight<br>Whole body fat mass | FTO | rs1421085 | intron |
| 17 | 7080316 | Alkaline phosphatase (quantile) | ASGR1 | rs55714927 | synonymous |
| 20 | 4157072 | Immature reticulocyte fraction | SMOX | rs6084653 | intron |
| 22 | 37462936 | Haemoglobin concentration | TMPRSS6 | rs855791 | missense |
| 22 | 44324727 | Alanine aminotransferase (quantile) | PNPLA3 | rs738409 | missense |
| 22 | 46364161 | 3mm strong meridian (left)<br>3mm weak meridian (left)<br>6mm strong meridian (left)<br>6mm weak meridian (left) | WNT7B | rs9330813 | intron |

For an **expanded version of this table**, which includes our explanation of the mechanism by which each SNP directly affects the trait(s) for which it an outlier, **see the attached xlsx file**.
